# Niche Overlap Determines Phage Predation Impact on Bacterial Diversity

**DOI:** 10.64898/2026.11.26.769424

**Authors:** Malick Ndiaye, Thania Sbaghdi, Lucie Kesner, Prajwal Padmanabha, Meline Garcia, Ena Bochaton, Sara Mitri, Philipp Engel

## Abstract

Bacteriophages kill a substantial fraction of the world’s bacteria every day, yet the effect of this mortality on bacterial community composition remains poorly understood. Here, we show that resource availability determines the ecological outcome of phage predation. We passaged six synthetic *Bifidobacterium* communities under simple or complex carbon conditions, with and without phages. Under simple nutritional conditions, phages drove kill-the-winner dynamics, suppressing dominant competitors and increasing diversity. Under complex nutritional conditions, niche partitioning limited competition and sustained stable bacteria-phage coexistence, with limited effect on diversity. A mathematical model confirmed that niche overlap among bacteria is sufficient to generate these contrasting outcomes. This dependence on the resource environment explains why the ecological effects of phages have proven difficult to generalize across ecosystems.

## Introduction

Ecological communities are structured by both “bottom-up” factors (e.g., resource availability, temperature, and pH) and “top-down” forces (e.g., predation and parasitism) (*1*–*3*). In the microbial world, top-down control is largely mediated by bacteriophages (hereafter, phages), which are a major source of bacterial mortality owing to their abundance and ubiquity across ecosystems worldwide (*4, 5*). These viruses have profoundly shaped bacterial evolution by mediating horizontal gene transfer and driving diversification through coevolution (*6*-*9*). Despite this clear evolutionary impact, their effects on community composition over ecological timescales remain poorly understood (*10*). Resolving this gap is important, as bacterial community composition is directly tied to key ecosystem functions, from biogeochemical cycling to host health (*1, 11, 12*). Moreover, emerging applications such as phage therapy and phage-mediated microbiome engineering ultimately depend on predicting the ecological impacts of phages on microbial communities (*13, 14*).

Among the main ecological effects attributed to phages is their potential to increase bacterial community diversity. In theory, by preying preferentially on abundant bacteria, phages suppress competitive taxa and promote coexistence (*15, 16*). However, support for these “kill-the-winner” dynamics from natural systems is mixed. Algal blooms collapse following peaks in viral abundance, resulting in increased community diversity (*17*-*19*). Yet this co-variation in densities is correlational and could equally reflect bottom-up effects (e.g., resource depletion) dampening bacterial growth, with phages tracking host dynamics rather than driving them (*10*). Across other ecosystems, phages seem to coexist stably with their hosts without driving pronounced compositional shifts, and environmental variables emerge as stronger determinants of community composition (*20*-*23*). Experimental work paints a similarly inconsistent picture: some studies report strong effects of phage addition on bacterial community composition (*24*-*26*), while others find little or no effect (*27*-*30*).

These contrasting observations suggest that the ecological impact of phages is strongly context-dependent. Variation in host density, diversity, and physiology is likely to modulate the ecological effects of phage predation. In turn, these community-level properties are strongly shaped by bottom-up environmental forces such as resource availability (*31*-*34*). Thus, a predictive understanding of phage ecology requires quantitatively linking phage effects to the environmental context that structures host communities.

The honeybee gut microbiome provides a tractable system for addressing this aim. It is dominated by a small number of conserved bacterial taxa that contribute to host health and are preyed upon by diverse phages (*35, 36*). Moreover, worker honeybees shift from a pollen-rich to a nectar-rich diet as they transition from nursing to foraging (*37*), providing a natural context for investigating how resource variation shapes the ecological impact of phage predation. Here, we combined growth experiments with mathematical modelling, using synthetic Bifidobacterium communities and their phages from the honeybee gut. By manipulating nutrient complexity and phage pressure, we investigated how bottom-up environmental conditions modulate the impact of phage predation on microbial community assembly. Our findings highlight the importance of the resource environment in shaping phage-bacteria interactions, helping explain why the ecological effects of phages can be difficult to generalize across ecosystems.

## Results

### Phage effects on alpha-diversity are contingent on resource complexity

We selected 12 *Bifidobacterium* strains representing the species- and strain-level diversity of this genus in the honeybee gut microbiome (fig. S1A; table S1). We then isolated phages that belonged to 30 viral operational taxonomic units (vOTUs; Methods; table S2) that collectively infected 11 of the 12 strains (See *Supplementary Results: Assembly of a Diverse Phage-Bacteria Synthetic Community that Recapitulates Ecological Survey*; fig. S1B-D). A pilot experiment showed that the phage-resistant strain (ESL0170) would dominate in the presence of phages (See *Supplementary Results: Phage effect is dampened by the dominance of a phage-resistant strain;* fig. S6). Thus, to prevent dynamics from being driven by this single resistant taxon rather than by phage-resource interactions, we excluded ESL0170 from subsequent experiments.

We divided the remaining 11 *Bifidobacterium* strains into six permuted communities (Bicom 1-6, biological replicates), each containing six strains (fig. S9). These communities were grown in a longitudinal, full-factorial experiment on a carbohydrate-free medium (cfMRS) supplemented with either sucrose water (SW, low-complexity resource) or pollen extract (PE, high-complexity resource), representing the two main dietary inputs of honeybees, i.e., nectar and pollen (*37*). We tested both resource conditions in the presence and absence of a mixture of isolated phages. Each community was transferred into fresh media every 48 hours for a total of 10 cultures (passages), with no new phages added (**Fig. 1A**; table S10). A phage contamination was detected in the no-phage control of Bicom2 in PE during passages 8-10, but a sensitivity analysis excluding these passages confirmed that this contamination did not affect the reported results (Methods; fig. S10).

**Figure 1:**
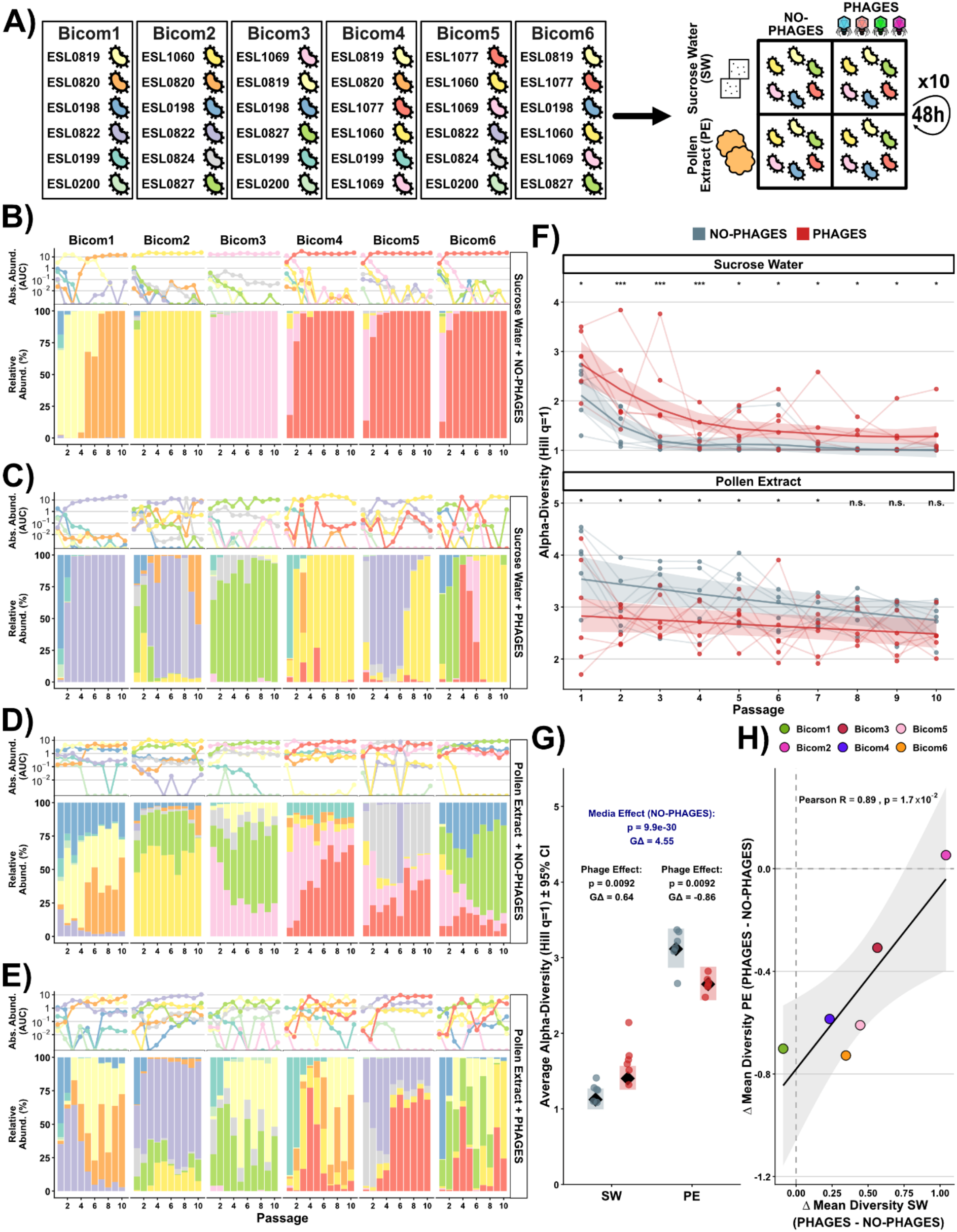
Phage effects on alpha-diversity are contingent on resource complexity. (**A**) Six synthetic communities (Bicom1-6), each composed of six Bifidobacterium strains inoculated at equal initial OD-based ratios (fig. S9), were grown in a full-factorial longitudinal experiment, in which two nutrient conditions, i.e., sucrose water (SW; low resource complexity) and pollen extract (PE; high resource complexity) were tested in the presence and absence of phages. Each Bicom in each treatment was serially transferred every 48h for a total of 10 cultures (passages). (**B-E**) Absolute abundances (top, expressed as OD-based Area Under the growth Curve, AUC) and relative abundances (bottom, stacked bar plot) of the strains in the six Bicoms across the 10 passages for each condition: (B) SW without phages, (C) SW with phages, (D) PE without phages, and (E) PE with phages. (**F**) Alpha-diversity (Hill number q=1) over time in SW (top) and PE (bottom) conditions, in the absence (grey) or presence (red) of phages. Connected points show individual community trajectories. Lines show generalized additive model (GAM)-predicted marginal means ± 95% CI. Asterisks denote significant pairwise differences between phage treatments at each passage (* p < 0.05, ** p < 0.01, *** p < 0.001; n.s., not significant). (**G**) Average alpha-diversity across passages summarized by condition. Diamonds show GAM’s estimated marginal means. Crossbars show 95% CI. Points show community averages across passages. Statistics indicate the effect of media (NO-PHAGES only) and the effect of phages within each media condition (GΔ, Glass’s Δ effect size). All p-values are FDR-corrected. (**H**) Correlation between phage-induced changes in mean alpha-diversity in SW (x-axis) and PE (y-axis) across communities. Each point represents one community (Bicom1-6). The line shows the linear regression fit ± 95% CI. Pearson R and p-value are indicated.

We first characterized community assembly in the absence of phages across the two resource conditions. In SW, we consistently observed competitive exclusion, with the strain exhibiting the highest productivity (AUC) in monoculture outcompeting all the others (**Fig. 1B**, fig. S7A; table S11). One exception occurred in Bicom1, where ESL0819 dominated during the first four passages before being overtaken by ESL0820, which can be explained by the unusually long lag-phase of ESL0820 (fig. S7B). In PE, a higher alpha-diversity than in SW was sustained at every passage, with strains distributed more evenly across the community (**Fig. 1D**). Quantified using effect sizes (Glass’s Δ; defined as the difference between means divided by the standard deviation of the reference group, here SW and/or NO-PHAGES), the effect of resource complexity on alpha-diversity was large (GΔ = 4.55, p < 0.001; Fig. 1G; table S12). Thus, our results suggest that increased resource complexity facilitates bacterial coexistence, most likely due to niche partitioning. This is consistent with empirical studies across diverse microbial communities (*31, 32*) and with resource-consumer models, in which the number of coexisting species is determined by the number of available resources (*34*).

The introduction of phages induced pronounced compositional shifts under both resource conditions, but their effect on alpha-diversity was more moderate and reversed in direction depending on the environment. In SW, phages suppressed the dominant competitors (**Fig. 1C**) and shifted community composition toward different dominant strains compared to no-phage communities, resulting in a significant increase in alpha-diversity over 10 passages (GΔ = 0.64, p=0.009; **Fig. 1F-G**). These observations are consistent with phage-mediated competitive release. Conversely, phage predation caused a net reduction in alpha-diversity in PE (GΔ = −0.86, p=0.009; **Fig. 1E-G**). Moreover, the magnitude of the phage effect depended on community composition: communities that experienced the largest diversity gains via competitive release in SW were the same communities most resilient to diversity loss in PE (Pearson R = 0.89, p = 1.7×10^−2^; **Fig. 1H**).

Together, these results show that phages influence community composition, but that their effects on diversity are context-dependent, with opposite effects in low- and high-complexity resource environments. Moreover, phage effects on diversity were much weaker than those imposed by bottom-up resource complexity.

### Phage predation drives temporal instability through rapid oscillations in high-competition environments

We next examined how phage predation and resource complexity shape the temporal dynamics and stability of the community. To do so, we quantified time-decay relationships by relating Bray-Curtis dissimilarity between samples to their temporal distance. We then fitted a linear model to each time-decay relationship (**Fig. 2A**), in which the intercept of the regression line at Δtime = 1 represents short-term turnover (compositional instability between two consecutive cultures; **Fig. 2B**), while the slope quantifies the rate of community divergence over time (**Fig. 2C**).

**Figure 2:**
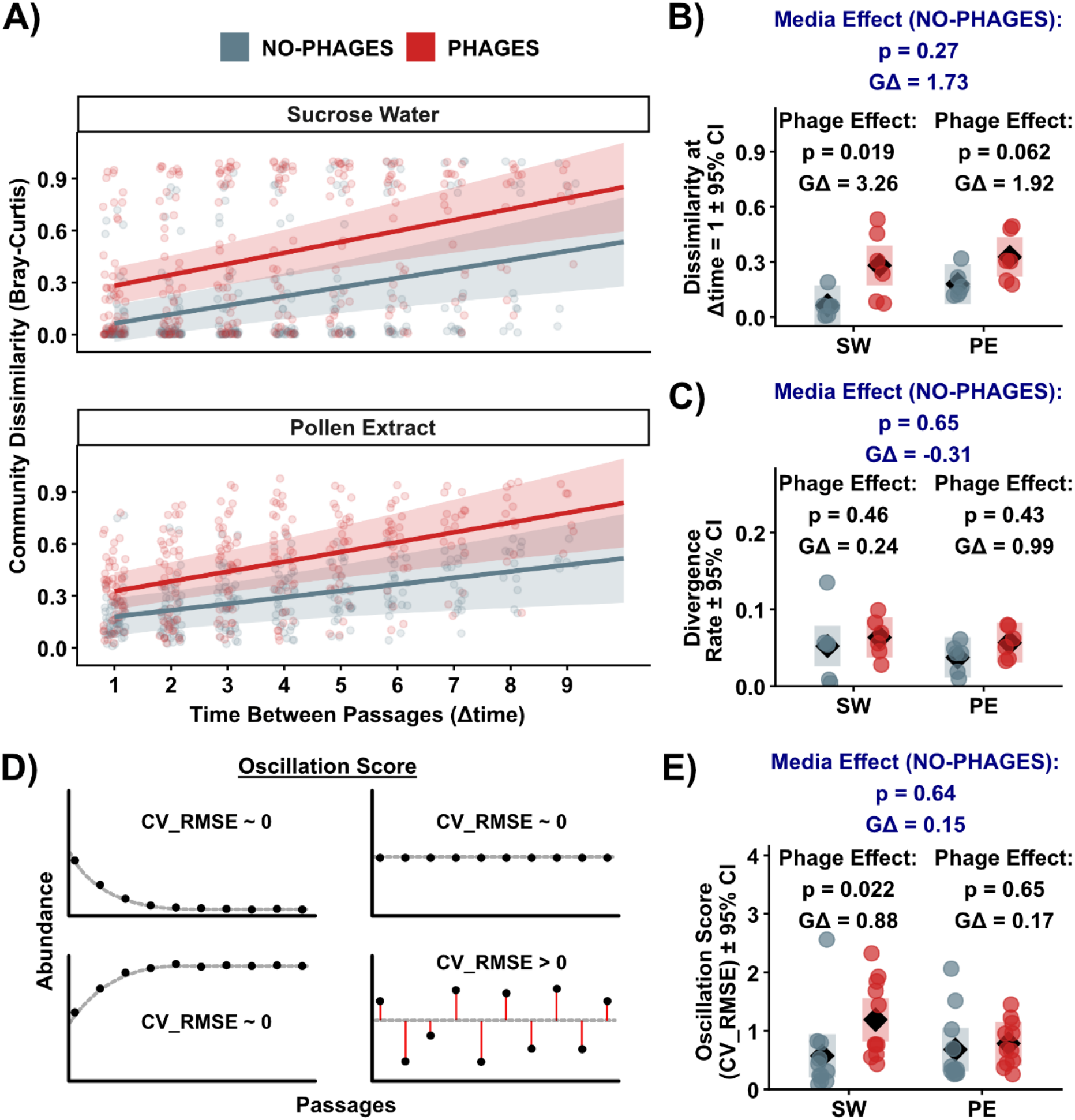
Phage predation increases community turnover and strain-level oscillations selectively under low resource complexity. (**A**) Time-decay relationship of community similarity in SW (sucrose water, top) and PE (pollen extract, bottom). Community dissimilarity (Bray-Curtis) is plotted against the number of passages separating any two timepoints (Δtime) for communities grown without (grey) or with (red) phages. Lines show linear regression fits ± 95% CI. (**B**) Short-term community turnover, measured as Bray-Curtis dissimilarity between consecutive passages (Δtime = 1), for SW and PE conditions. Diamonds show estimated marginal means. Crossbars show 95% CI. Points show community-level averages. (**C**) Community divergence rate, measured as the slope of the Bray-Curtis dissimilarity over Δtime, for SW and PE conditions. Diamonds, crossbars and points as in (B). (**D**) Schematic illustration of the oscillation score (CV_RMSE). Strains reaching carrying capacity, going extinct, or remaining at steady state yield an oscillation score near zero, whereas strains exhibiting oscillatory dynamics around a mean abundance yield an oscillation score greater than zero. Red lines indicate residuals from the best-fitting non-oscillating model. (**E**) Oscillation scores of individual strains, averaged per community, for SW and PE conditions. Diamonds, crossbars and points as in (B). For all crossbar plots, statistics indicate the effect of media (NO-PHAGES only) and the effect of phages within each resource condition (GΔ, Glass’s Δ effect size). All p-values are adjusted for multiple comparisons using the FDR method.

Our analysis revealed that resource complexity had no effect on short-term turnover, whereas phage predation increased it, as reflected by higher Bray-Curtis distances at Δtime = 1 (**Fig. 2B**; table S13). This effect was more pronounced and statistically significant in SW (GΔ = 3.26, p = 0.019) than in PE (GΔ = 1.92, p = 0.062), indicating that simple resource conditions exacerbate the compositional variability induced by top-down pressure. Notably, neither phage predation nor resource complexity significantly altered the slope of the time-decay relationship (**Fig. 2C**). Thus, although phages increased passage-to-passage compositional turnover, they did not accelerate long-term community divergence.

We then asked whether this short-term compositional variability translated to functional instability at the community level. Using longitudinal OD measurements across passages (fig. S11A-B), we calculated the area under the growth curve (AUC) for each community at each passage as a proxy for productivity (fig. S12A). Phage-treated communities exhibited a significantly higher coefficient of variation (CV) in AUC than NO-PHAGES controls, indicating greater temporal variability in productivity (fig. S12B; table S14). Together, these results show that phages destabilize both community composition and the consistency of community-level growth over time.

The combination of high short-term community turnover and variation in biomass, without a corresponding increase in long-term community divergence, suggests that individual strains undergo oscillatory dynamics over time. To quantify this, we calculated an “oscillation score” (*CV_RMSE*, see Methods) which captures how much a strain’s absolute abundance fluctuates over time relative to three models of non-oscillating growth: reaching carrying capacity, going extinct, or remaining at steady state (CV_RMSE∼0), with higher scores indicating greater fluctuation (**Fig. 2D**; fig. S13A-C). Consistent with the beta-diversity results, bacterial strains exhibited significantly higher oscillation scores (*CV_RMSE>0)* in the presence of phages only in SW (**Fig. 2E**; table S14).

Collectively, these findings demonstrate that top-down predation is the primary driver of temporal instability in these synthetic communities. This instability arises from rapid fluctuations in strain abundances and is amplified under strong resource competition, suggesting that niche partitioning in complex resource environments can dampen predator-induced boom-and-bust dynamics.

### Phage-mediated expansion of the effective species pool occurs only under high-competition conditions

To estimate the effective species pool maintained over the course of the experiment, we integrated within-timepoint diversity (alpha-diversity) and turnover between timepoints (beta-diversity) by calculating temporal gamma-diversity for each community (**Fig. 3**; Methods). In SW, phage predation increased the effective species pool (GΔ = 2.72, p = 0.021), indicating that more taxa persisted over time. In PE, resource complexity alone supported substantially higher baseline diversity (GΔ = 4.58, p = 0.00038), but phage addition had no significant effect on the effective species pool (GΔ = 0.92, p = 0.36). This absence of a phage effect on gamma diversity in PE is consistent with their negative impact on alpha-diversity being offset by a positive effect on beta diversity. Thus, phage-mediated expansion of the effective species pool is restricted to simple resource environments (Table S15).

**Figure 3:**
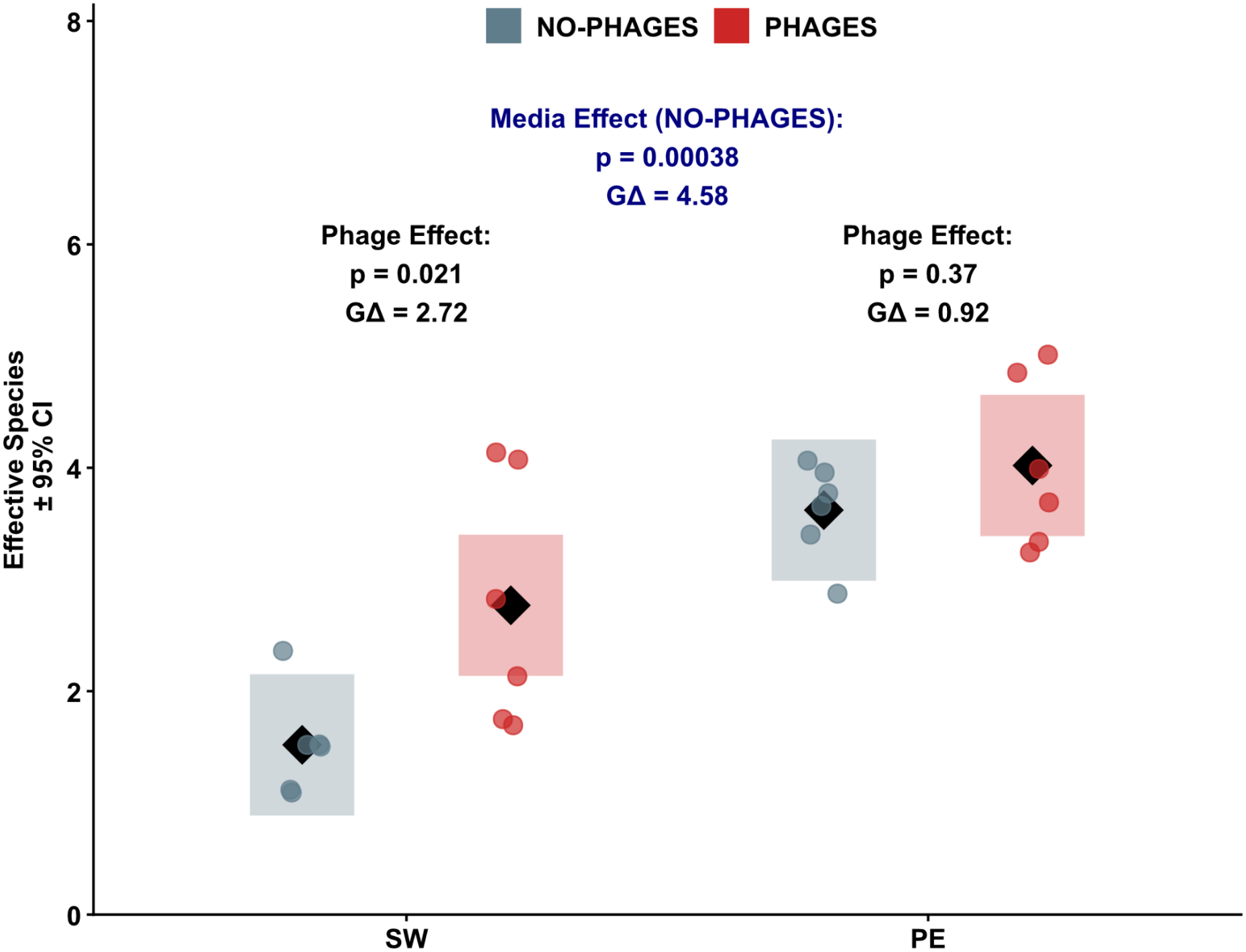
Phage predation expands the effective species pool only under low resource complexity. Temporal gamma-diversity provides an estimate of the effective species pool maintained across the ten passages. Bars show estimated marginal means ± 95% CI from a linear mixed model (Media × treatment, with community as random effect). Points show individual community values. Statistics indicate the effect of media (NO-PHAGES only) and the effect of phages within each media condition. Effect sizes are reported as Glass’s Δ. All p-values are adjusted for multiple comparisons using FDR correction. SW, sucrose water; PE, pollen extract.

### Resource complexity sustains viral phenotypic diversity and stability

To investigate the mechanisms underlying phage-driven changes in diversity, we longitudinally tracked viral titers of phages targeting ESL1077 and ESL1060, two strains suppressed upon phage addition despite being dominant in their absence (**Fig. 1B-E**). Both strain-phage pairs exhibited sporadic predator-prey oscillations, although these dynamics were poorly synchronized. Phage titers occasionally peaked alongside increases in bacterial densities, but phage populations frequently declined to undetectable levels and did not consistently track bacterial dynamics (fig. S14A-D). Since these phages infect multiple hosts (fig. S1C, E), we reasoned that their population dynamics are shaped by multiple community members rather than any single strain. Thus, we turned to a community-level characterization of phage infectivity to better resolve how resource complexity shapes predator-prey dynamics.

To this end, we performed longitudinal spot assays against all 11 *Bifidobacterium* strains to generate a community-level infectivity profile across the ten passages (fig. S15; table S16). We qualitatively scored plaque formation from 0 (no plaque) to 3 (strong plaque) to generate an “infectivity profile” for each viral community across the ten passages (fig S15A-B). This assay revealed that phage communities maintained significantly higher plaque richness (i.e., the number of strains showing plaques at a given passage) in PE than in SW (GΔ = 0.93, p = 0.031; Fig. 4A-B), suggesting that the increased niche space maintained by resource partitioning in PE sustains a more phenotypically diverse phage population.

**Figure 4:**
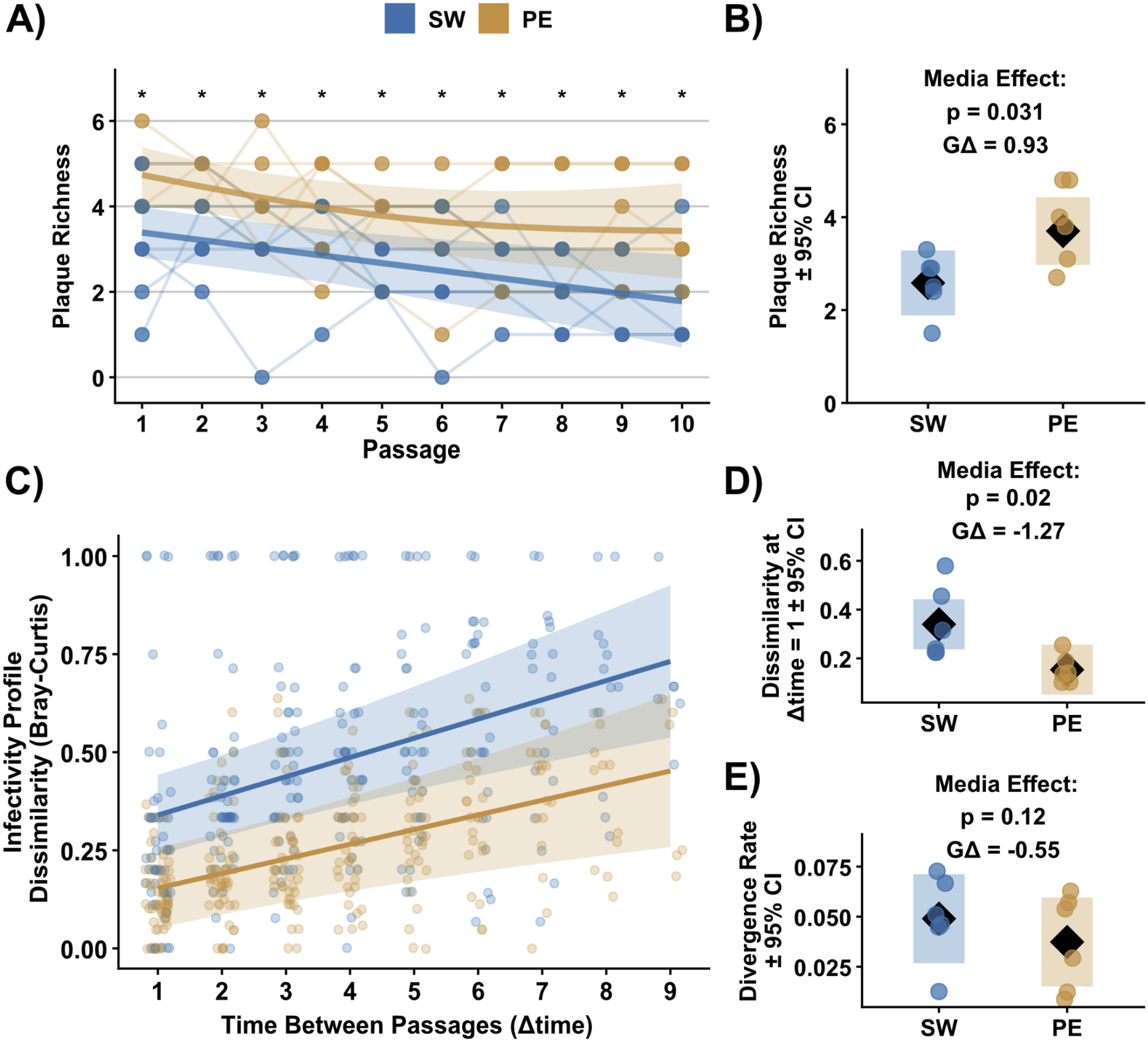
Resource complexity sustains phage functional diversity and stabilizes infectivity dynamics. (**A**) Plaque richness (i.e., the number of bacterial strains successfully lysed in each community) over ten passages for phage communities grown in sucrose water (SW, blue) and pollen extract (PE, orange). Connected points show individual community trajectories. Lines show generalized additive model (GAM) fits ± 95% CI. Asterisks indicate significant pairwise differences between media at each passage (FDR-corrected; * p < 0.05). (**B**) Average plaque richness summarized across passages. Diamonds show estimated marginal means, while crossbars show 95% CI from the GAM. Points show community-level averages. Effect size is reported as Glass’s Δ. (**C**) Time-distance decay relationship of the infectivity profiles. Bray-Curtis dissimilarity between infectivity profiles is plotted against the number of passages separating any two timepoints (Δtime) for SW and PE communities. Lines show linear regression fits ± 95% CI. (**D-E**) Short-term turnover of infectivity profiles, measured as Bray-Curtis dissimilarity between consecutive passages (Δtime = 1; D), and turnover rate, measured as the slope of dissimilarity over Δtime (E). Diamonds show estimated marginal means, while crossbars show 95% CI from linear mixed-effect model. Points show community-level values. Effect sizes are reported as Glass’s Δ.

Infectivity profiles were also more temporally stable in PE, with SW communities displaying higher short-term turnover (**Fig. 4C-E**; table S17). Together, these results show that bottom-up effects of resource complexity propagate up the trophic levels to shape the composition and dynamics of the phage population. Moreover, the higher turnover of infectivity profiles in SW mirrors the bacterial oscillations observed in the host community (**Fig. 2E**). This suggests that predator-prey dynamics, while difficult to resolve at the level of individual strain-phage pairs, are detectable across the community as a whole. Consistent with this, predator-prey dynamics were more synchronized in our pilot experiment in SW (fig. S16; See *Supplementary Results: Phage effect is dampened by the dominance of a phage-resistant strain*). In that experiment, community dynamics were effectively dominated by the interplay between a more competitive but susceptible strain (ESL1077) and a phage-resistant strain (ESL0170), reducing the system to a near two-strain interaction (fig. S6B, S16).

### A mathematical model links niche partitioning to the context-dependent effects of phage predation

To assess whether our findings generalize beyond the communities studied here, we developed a mathematical model of bacterial community dynamics under phage predation and resource competition (**Fig. 5A**; Methods; see Model Description). Phage-host interactions were assigned at random, and the SW condition was modeled as bacteria accessing only a subset of the resources available in PE.

**Figure 5:**
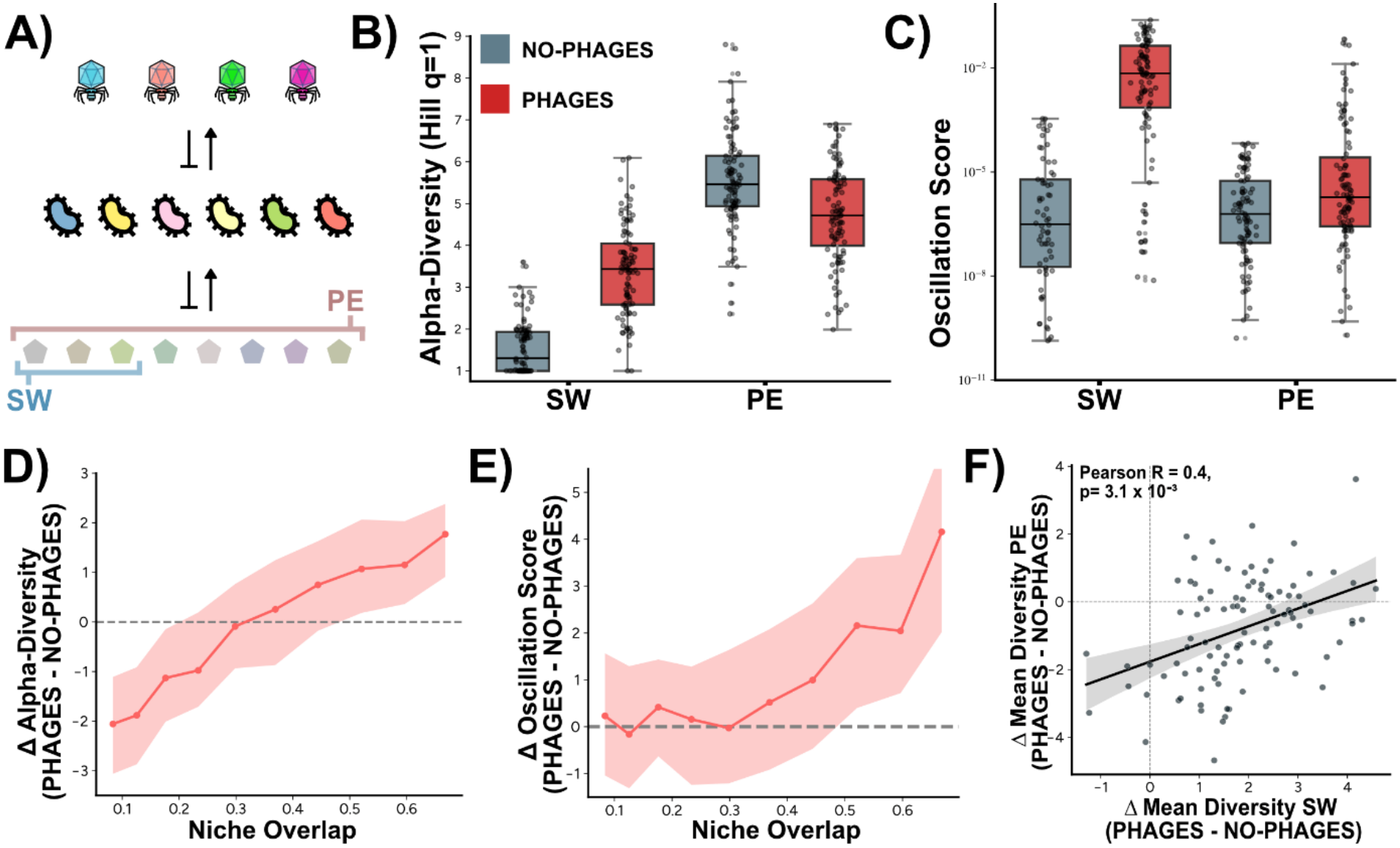
A mathematical model recapitulates experimental findings and identifies niche overlap as sufficient to explain the context-dependence of phage predation effects. (**A**) Model schematic. Phages (top) interact with a community of bacterial strains (middle), which are competing for a shared resource pool (bottom). The sucrose water (SW) condition is modeled as bacteria accessing a limited subset of available resources, while pollen extract (PE) provides the full resource pool. (**B-C**) Model output for (B) alpha-diversity (Hill number q=1) and (C) oscillation score across simulated communities grown with (red) or without (gray) phages under SW and PE conditions. Each point represents one simulated community (n=100). (**D-E**) Effect of niche overlap on phage-induced changes in (D) alpha-diversity and (E) oscillation score (log10), expressed as the difference between phage and no-phage conditions (PHAGES − NO-PHAGES). Lines show mean effects ± 25-75 quantiles across simulations (n=50 per niche overlap value). (**F**) Correlation between the phage-induced change in mean alpha-diversity in SW and PE across simulated communities. Each point represents one simulated community. The line shows the linear regression fit ±95 % CI. Pearson R and p-value are indicated in the plot.

The model qualitatively reproduced our experimental results. Phage predation increased alpha-diversity in SW while decreasing it in PE, and oscillation scores rose in the presence of phages only in SW (**Fig. 5B-C**). In the model, niche overlap is determined by the similarity of strain growth rates across resources in the strain-resource matrix, with higher overlap indicating that two strains grow similarly on the same resource. Increasing niche overlap reversed the effect of phage predation on alpha-diversity (**Fig. 5D**) and led to higher oscillation scores in the presence of phages (**Fig. 5E**). These results were robust to variation in resource number, trade-offs between growth and phage susceptibility, burst size, trait scales, and total carbon supply (fig. S17A-B). Thus, niche overlap among community members alone is sufficient to explain why the ecological effects of phage predation depend on resource complexity.

Finally, the model reproduced the correlation observed experimentally between phage effects on alpha-diversity across resource conditions (**Fig. 1H**): communities that gained more alpha-diversity in response to phages in SW tended to lose less alpha-diversity in PE (**Fig. 5F**). This correlation was significant in all 24 parameter configurations tested (fig. S17C), suggesting that it reflects a robust property of phage-bacteria interactions and that the ecological impact of phages in one resource context can predict their impact in another.

## Discussion

A central question in microbial ecology is the relative contribution of bottom-up and top-down forces to community assembly. Our results show that these two ecological forces are not independent. The resource environment is the major determinant of bacterial community diversity, while also shaping the extent to which phage predation can influence that diversity. Under simple resource conditions, intense competition creates the conditions for kill-the-winner dynamics, enabling phages to promote diversity through competitive release. Under complex resource conditions, strains partition niches rather than competing directly, limiting how much phage predation can increase diversity. This dependence on environmental context may help reconcile contrasting reports on the ecological effects of phage predation across systems, and may explain why these effects have proven so difficult to generalize (*10, 38*). Strong top-down effects are most often detected in simple or competitive systems where phages are the dominant ecological pressure (*24*-*26*), such as small synthetic communities (*24*) or algal blooms (*17*-*19*). In contrast, these effects weaken or disappear in complex communities where phages are only one of many interacting ecological drivers (*27*-*30*). For instance, reintroducing fecal-derived phages into a complex bacterial community cultured in fiber-rich media had minimal effect on overall composition (*30*), consistent with our finding that resource complexity buffers communities against phage-driven diversity shifts. Rather than asking whether phages drive community diversity, our results suggest the more productive question is under what conditions they do so.

Previous explanations for the context-dependent effects of phages on microbial communities have invoked shifts in phage lifestyle, nutrient limitation, or changes in host physiology (*4, 39, 40*). For example, the piggyback-the-winner hypothesis proposes that high bacterial densities favor lysogeny over lytic replication, reducing phage impacts on community composition (*41*). Resource availability itself can also affect host susceptibility to phages, potentially shifting their ecological impact across nutrient environments (*42, 43*). Our model, however, reproduces the same qualitative pattern through niche partitioning alone, without changing phage lifestyles or host physiology. When hosts occupy distinct niches, no single strain dominates, limiting the conditions for boom-and-bust dynamics and yielding more stable phage-host coexistence over time. The model also recapitulated a finer-grained result: communities that gained the most diversity from competitive release in SW were the same communities most resilient to diversity loss in PE. The mechanism behind this correlation remains unclear. One possibility is that phage predation pushes each community toward a similar diversity steady state, regardless of resource environment: communities with a higher diversity steady state would gain more in SW and lose less in PE. In turn, this steady state should be predictable from the host-phage infectivity matrix. However, testing this would require a larger panel of communities than examined here. Still, our results suggest that niche overlap alone can predict both the direction and magnitude of phage effects across resource environments, offering a parsimonious explanation for how diverse bacterial and phage communities coexist in nature (*21, 22*).

In pollen extract, phages had a limited effect on community diversity, yet they altered both community biomass and composition. This suggests that even in less competitive environments, phages can exert strong selective pressures during the early stages of community assembly, thereby shaping downstream community structure and function without necessarily changing overall diversity. For instance, lytic phage predation has been proposed to modulate the assembly of the human gut microbiome during early infancy (*44*).

There is growing interest in harnessing phages for microbiome engineering and treating antibiotic-resistant infections. Our findings show that both the magnitude and nature of phage effects on bacterial communities depend strongly on the bottom-up forces that structure the host community, providing a more predictive framework for understanding phage impacts. For example, phage-based treatments for gastrointestinal diseases are of increasing interest (*45*), and our results suggest that patient diet could influence treatment outcomes. Here, we focused on a single bottom-up variable: resource complexity. Extending this work to other environmental factors (e.g., temperature, O_2_ levels) will be essential to assess the generalizability of our results. Finally, the phage-bacteria synthetic community established in this study should be amenable to *in vivo* experiments in honeybees. Such work would help clarify whether dynamics observed in our homogeneous, well-mixed *in vitro* system persist in more complex settings where host-associated factors (e.g., spatial structure and immune responses) may further modulate phage-bacteria interactions.

## Acknowledgments

We thank Julien Marquis, Aline Charpagne, Melanie Dupasquier, Corinne Peter, and Johann Weber from the Lausanne Genomic Technologies Facility (GTF) for technical support during DNA extraction, library preparation, and sequencing. Anthropic’s Claude was used to edit and polish this manuscript.

## Author contributions

Conceptualization: M.N., L.K., and P.E.; Methodology: M.N., T.S., and L.K.; Formal analysis: M.N.; Data curation: M.N., T.S., L.K.; Investigation: M.N., T.S., L.K., M.G., and E.B.; Mathematical Modeling: P.P. and S.M.; Writing - original draft: M.N., and P.E.; Editing: M.N., T.S., P.P., S.M., L.K., M.G., and P.E.; Resources: P.E.; Project administration: P.E.; Funding acquisition: P.E.

## Data availability

Raw sequencing data (shotgun metagenomic and full-length 16S rRNA gene amplicon) and the newly sequenced *Bifidobacterium* genomes have been deposited in the NCBI Sequence Read Archive under BioProject PRJNA1484827, with BioSample accessions SAMN61269747 to SAMN61269767; table S1 gives the strain-to-accession mapping. Bacterial isolate genomes, viral MAGs, intermediate data files, and the tables and scripts used to generate the figures are archived at Zenodo https://doi.org/10.5281/zenodo.23616972. Analysis code is available at https://github.com/MalickNdiye/PHIFIDO and archived at Zenodo https://doi.org/10.5281/zenodo.22759207. All *Bifidobacterium* strains generated in this study are available from the corresponding author upon request. The enriched phage cocktails exist as finite lysate stocks and cannot be shared in their original composition. Nonetheless, the enrichment procedure for deriving equivalent cocktails starting from honeybee gut homogenates is described in the Materials and Methods.

## Supplementary Materials list

Supplementary Results

Materials and Methods

Model Description

Figures S1 to S18

Tables S1 to S25

