## Supplementary Materials for "Niche Overlap Determines Phage Predation Impact on Bacterial Diversity"

#### Assembly of a Diverse Phage-Bacteria Synthetic Community that Recapitulates Ecological Survey

Our study aimed to assemble a synthetic phage-bacteria community that simulates a natural one. Thus, we focused on isolating a diverse community of phages and bacteria that coexist in the same environment. Our prior ecological survey of the honeybee gut microbiome revealed modular phage-bacteria interactions, where phage and bacterial diversity and composition correlated within the same module *(35)*. One such module encompassed all *Bifidobacterium* species found in the honeybee gut. To replicate this in our synthetic community, we selected 12 *Bifidobacterium* strains (fig. S1A) that represent a broad diversity of *Bifidobacteria* found in the honeybee gut microbiome. These strains belonged to 6 out of 10 distinct *Bifidobacterium* species in our genome collection.

Next, we adopted an enrichment approach to isolate phages that target the bacterial members of our synthetic community (fig. S1B, fig. S2). Three gut homogenates, each obtained by pooling guts from 170-120 bees, were enriched for VLPs using a series of centrifugations and filtrations (Methods), yielding three phage cocktails (hereafter, cocktails A, B, and C; table S3). These initial cocktails were then incubated with each individual *Bifidobacterium* strain in the synthetic community, as well as with a mix of all twelve strains in sucrose water (SW) or pollen extract (PE). This incubation lasted for 48 hours, after which the phages were recovered from the culture by filtration and re-incubated with the ancestral bacteria for two additional passages. Alongside these cultures, a control without phage addition was run for each strain or strain mix. The aim of this approach was to enrich for phages capable of replicating in a single strain or in a mix of all *Bifidobacterium* strains, while depleting those unable to do so.

Following phage enrichment across three successive cultures, we obtained 56 phage cocktails (3 original cocktails × (12 strains + 2 strain mixes) + 14 controls). To determine which phages were enriched on each strain, these cocktails were subjected to metagenomic sequencing (fig. S1C; table S2). The resulting phage contigs were clustered at 95% average nucleotide identity and 85% alignment fraction to define viral Operational Taxonomic Units (vOTUs). In total, we enriched 30 medium- to high-quality vOTUs (Methods) targeting all bacterial strains except ESL0170 and ESL0824 (table S4). Of these, 29 belonged to the class *Caudoviricetes* and one to *Malgrandaviricetes* (table S2). Among the recovered vOTUs, nine were predicted to be temperate, whereas the remainder were lytic (table S2). Each enriched cocktail contained a median of two vOTUs, but dominance within cocktails was high (median dominance = 0.72), with a single vOTU dominating cocktails enriched on most individual strains (fig. S1D). Notably, cocktails enriched on the 12-strain mixes contained subsets of the vOTUs recovered from single-strain enrichments (fig. S1C).

The bacterial strain used for enrichment explained substantially more variation in vOTU composition (PERMANOVA, $\omega^2 = 0.37$, $p = 0.0001$) than the cocktail of origin ($\omega^2 = 0.04$, $p = 0.007$), indicating strain specificity of the enriched phages (table S5). Of the enriched vOTUs, 27 out of 30 were predicted to target *Bifidobacterium* based on CRISPR spacer-protospacer matches (fig. S3A; table S6). Moreover, most *Bifidobacterium* vOTUs present in the initial phage cocktails were also recovered after enrichment (fig. S3B), indicating that the enriched cocktails capture a large fraction of the *Bifidobacterium* phage diversity present in natural honeybee populations. Finally, although four vOTUs were detected in control samples, none formed plaques on any of the 12 strains (fig. S4A). Further analysis revealed that three of these vOTUs correspond to constitutively expressed prophages from cultured strains (ESL0170, ESL0198, ESL0819), while one vOTU likely represented contamination during DNA extraction (fig. S4B-D).

We conducted plaque assays for all possible combinations of enriched cocktails and bacterial strains (fig. S1E; table S4). All strains except ESL0170 were lysed by at least one of the enriched cocktails. Those derived from homogenates B and C showed similar lytic activity profiles across strains, whereas cocktails derived from initial cocktail A exhibited reduced activity that largely overlapped with the host range observed for B- and C-derived cocktails. Most enriched cocktails infected multiple strains and displayed broad host ranges, often extending to phylogenetically distant species, while failing to infect some closely related strains. Consistently, we found no correlation between host phylogenetic distance and either cocktail lytic activity or enriched phage community composition (fig. S5A). Moreover, vOTU composition alone poorly predicted plaque formation, suggesting that cocktails containing the same vOTU can differ in lytic activity (fig. S5B). This pattern likely reflects within-vOTU genetic variation affecting phage host range.

In conclusion, we established a diverse synthetic bacterial community of *Bifidobacterium* targeted by a diverse set of phages with overlapping host ranges.

#### Phage effect is dampened by the dominance of a phage-resistant strain

To investigate the relative contributions of top-down predation and bottom-up resource complexity, we first established a synthetic community comprising all 12 *Bifidobacterium* strains. This community was subjected to a longitudinal, full-factorial experiment in which it was incubated in carbohydrate-free MRS supplemented with either sucrose water (SW) or pollen extract (PE) medium, with or without a cocktail containing all B- or C-derived phages. Each condition had 4 technical replicates that were serially passaged every 48 hours for 15 transfers (fig. S6A; table S7). As detailed below, this experiment showed that a single phage-resistant strain (ESL0170) dominated the community under phage predation in both environments. Thus, we excluded it from all subsequent experiments.

In the absence of phages (fig. S6B), community assembly in SW was dominated by the most productive strain in monoculture (as assessed by the growth Area Under the Curve, AUC), ESL1077, consistent with the competitive exclusion principle (fig. S7A). Phage addition suppressed ESL1077 but consistently promoted the dominance of the phage-resistant strain ESL0170, which rapidly outcompeted all other strains (fig. S6B). Because a single strain dominated with or without phages, alpha-diversity was unaffected, except for a few passages in the phage condition where ESL1077 rebounded in relative abundance, producing alpha-diversity peaks (fig. S6D-E; table S8). A similar pattern emerged in PE; while niche partitioning allowed for high coexistence in the absence of phages (fig. S6C), the presence of predators again led to a decrease in diversity due to the dominance of the phage-resistant ESL0170 (fig. S6D-E; table S8).

Alongside community composition, we quantified total community productivity by tracking longitudinal optical density (OD) and calculating growth area under the curve (AUC) across passages (fig S8A-B). This analysis revealed that phage predation consistently destabilized community-level functional output. In SW, productivity declined sharply, reaching a minimum around passage 5, then gradually recovered. In PE, phage presence induced rapid oscillations in AUC around the NO-PHAGE baseline. Consequently, in both environments, communities exposed to predation exhibited significantly greater variability in total productivity (higher AUC coefficient of variation) compared to their respective controls (fig S8C; table S9).

Together, these results demonstrate that phages strongly impact both community composition and function. Moreover, C- and B-derived phages were indistinguishable in their effects on bacterial communities. However, in this 12-strain system, the dynamics converged on a predictable outcome driven by phage immunity rather than by the interaction between phage predation and resource complexity. In both environments, community assembly collapsed toward dominance by the resistant strain ESL0170. Although the resulting functional instability confirms sustained top-down pressure, this resistance-driven outcome obscures the interplay between phage predation and bottom-up resource effects on community dynamics. While the presence of a fully phage-resistant strain is unlikely in natural communities, this result provides insight into how the emergence of resistance can rapidly restructure community assembly. Ultimately, these results motivated the exclusion of ESL0170 to better control community composition in subsequent 6-strain permuted experiments and to restrict phage selection for the more diverse cocktail B.

### Niche Overlap Determines Phage Predation Impact on Bacterial Diversity - Materials and Methods

#### Bifidobacterium Strains Isolation & DNA Extraction

To complement our pre-existing collection of *Bifidobacterium* strains from the honeybee gut, we isolated 27 additional strains (ESL1057 - 1088; table S1). Honeybee gut homogenates were serially diluted and plated onto pre-reduced MRSA agar plates (DSMZ 11b medium without glucose, supplemented with 2% fructose and 0.05% L-cysteine; pH 6.2-6.4). Plates were incubated anaerobically at 35°C for 3 days.

Single colonies were picked and resuspended in 160 µL of Buffer P1 (Qiagen, #1014630) for high-molecular-weight DNA extraction. Cells were lysed by adding 20 µL of lysozyme (100 mg/mL) and then incubated at 37°C with shaking at 600 rpm for 1 h. Subsequently, 20 µL of Proteinase K (20 mg/mL; Roche, #03115852001) was added, followed by incubation at 56°C with shaking at 600 rpm for 30 min. Next, 150 µL of Buffer AL (Qiagen, #19075) was added, and samples were incubated at 37°C with shaking at 600 rpm for 30 min. Cellular debris were pelleted by centrifugation at 14,000 rpm, and the resulting supernatant was transferred to a new tube containing 35 µL of 3 M sodium acetate and 270 µL of isopropanol for DNA precipitation at 4°C for 1 h. DNA was pelleted by centrifugation at maximum speed for 10 min at 25°C. The pellet was washed with 1 mL of ethanol, air-dried, and resuspended in 50 µL of TE buffer.

DNA was further purified using 40 µL of NGSClean magnetic beads (LabGene, #CNGS0050) with 5 min incubation at room temperature. The supernatant was discarded, and the beads were washed twice with 200 µL of 80% ethanol. DNA was eluted from the beads in 100 µL of 5 mM Tris-HCl (pH 8.0).

DNA integrity was assessed by electrophoresis. DNA concentration was measured using a Qubit fluorometer.

#### Whole Genome Sequencing of Bifidobacterium Strains

Genomic DNA from the newly isolated *Bifidobacterium* strains (ESL1057-ESL1088) was sequenced at the Lausanne Genomic Technologies Facility (GTF) using the Pacific Biosciences Sequel II platform to generate high-fidelity (HiFi) long reads. Genome assemblies were performed at the GTF using SMRTLink v11 (Pacific Biosciences) with default parameters. The resulting assemblies were used for all downstream analyses.

#### Growth Media Preparation

All experiments requiring bacterial growth were conducted in carbon-free MRS (cfMRS) medium supplemented with either 1% (v/v) sucrose water (SW; prepared from a 50% [w/v] sucrose stock solution) or 10% (v/v) pollen extract as the sole carbon source. Detailed information on the preparation of pollen extract is provided in the *Supporting Methods* section of Kešnerová et al. (2017) (*46*).

To prepare 1 L of cfMRS medium, the following components were mixed together: 10 g Bacto Peptone (Gibco, #211677), 5 g yeast extract (Oxoid, #LP0021B), 5 g sodium acetate (Sigma, #S2889), 2 g tri-ammonium citrate anhydrous (Fluka, #09723), 2 g dipotassium hydrogen phosphate (VWR, #26930.293), 0.2 g magnesium sulfate heptahydrate (Sigma), 0.05 g manganese sulfate monohydrate (Fluka), and 1 mL Tween 80. The volume was adjusted to 1 L with distilled water, and the pH was adjusted to 6.3-6.5 using HCl. The medium was autoclaved prior to use.

For bacterial pre-cultures, cfMRS was supplemented with 0.05% (w/v) sterile-filtered L-cysteine-HCl (Merck, #30120) together with the appropriate carbon source.

For all experiments involving phages (including no-phage controls), 2% (w/v) polyvinylpyrrolidone (PVP; Sigma, #P5288) was added to the cfMRS medium after pH adjustment and prior to autoclaving. Following sterilization, the medium was supplemented with sterile-filtered $CaCl_2$ (final concentration 25 mM), $MgCl_2$ (final concentration 100 mM), and 0.05% (w/v) L-cysteine-HCl, in addition to the required carbon source.

**Bacterial Growth Conditions**

*Bifidobacterium* strains were first plated from glycerol stocks on MRS agar supplemented with 2% fructose and 0.1% L-cysteine. Plates were incubated anaerobically at 35°C for 2-3 days. To ensure culture purity, single colonies were re-streaked onto fresh MRS agar plates and incubated under the same conditions for an additional 2-3 days.

Following the second incubation, cells were harvested from the plates and resuspended in 1 mL of 1× PBS in tubes containing 0.75-1 mm glass beads (Carl Roth). To prevent cell clumping, suspensions were homogenized using a FastPrep-24 5G homogenizer (MP Biomedicals) at 6.0 m/s for 15 s.

Cell suspensions were then diluted to an OD600 of 0.1 in 7 mL cfMRS supplemented with sucrose water to initiate pre-cultures. To ensure that all strains reached at least mid-exponential phase prior to harvesting, slow-growing strains (ESL0198, ESL0199, ESL0200, ESL0819, ESL0820, ESL0822, ESL0824) were incubated for 44-48 h, whereas fast-growing strains (ESL0170, ESL0827, ESL1060, ESL1069, ESL1077) were incubated for 20-24 h.

After pre-culturing, cells were pelleted by centrifugation at 6,000 × g for 2 min. The supernatant was discarded, and the pellets were resuspended in 1 mL PBS. Suspensions were adjusted to an OD600 of 0.5 in PBS and used to inoculate experimental cultures to a final OD600 of 0.05. Experimental growth was performed at 35°C under 5% $CO_2$.

For the synthetic phage-bacteria community passaging experiment involving six bacterial communities of different composition, a modified preparation protocol was used for practicality. Strains were pre-cultured as described above. After pre-culturing, cells were pelleted by centrifugation at 6,000 × g for 2 min and resuspended in PBS. Concentrated suspensions were mixed with glycerol to a final concentration of 20% (v/v) and adjusted to $OD_{600}$ = 10 before storage at -80°C as glycerol stocks in aliquots of

50 µL. Aliquots were thawed and resuspended directly in the appropriate medium to initiate experiments.

**Phage Cocktail Preparation from Conventional Honeybees**

For each initial phage cocktail (A, B, and C), 170-120 worker bees of *Apis mellifera carnica* (nurses and foragers) were collected from hives located at the University of Lausanne (Switzerland; table S3). Hindguts were dissected from individual bees and pooled in a 50 mL Falcon tube containing a 1:1 mixture of 0.75-1 mm glass beads (Carl Roth, #A554.1) and 0.1 mm zirconia/silica beads, together with 30 mL of SM buffer.

SM buffer was prepared as follows (per liter): 5.8 g NaCl, 2 g $MgSO_4 \cdot 7H_2O$, and 50 mL of 1 M Tris-HCl (pH 7.5), with the volume adjusted to 1 L using distilled water, followed by autoclaving.

Gut samples were homogenized using a FastPrep-24 5G homogenizer (MP Biomedicals) at 6.0 m/s for 45 s. Large debris was removed by centrifugation at 10,000 × g for 10 min at 4°C. The supernatant was sequentially filtered through 0.45 µm and 0.22 µm pore-size cellulose acetate filters (Clearline #037045 and Clearline #037044) to remove residual bacterial cells and debris. The resulting filtrate constituted the initial phage cocktail.

**Phage Enrichment Experiment**

Individual strains (ESL0170, ESL0198, ESL0199, ESL0200, ESL0819, ESL0820, ESL0822, ESL0824, ESL0827, ESL1060, ESL1069, ESL1077) pre-cultured in cfMRS supplemented with sucrose water were adjusted to an $OD_{600}$ of 0.5 in PBS. An 800 µL aliquot from each strain was combined to generate a 12-strain synthetic community mix.

Each individual strain and the mixed community were inoculated into 10 mL of cfMRS supplemented with either sucrose water (SW) or, for the community only, pollen extract (PE), at a final $OD_{600}$ of 0.05. Subsequently, 500 µL of the corresponding phage cocktail was added to each culture. Control cultures received an equivalent volume of SM buffer instead of phage suspension.

To monitor growth dynamics, a 150 µL aliquot from each culture was transferred to a 96-well plate for OD measurements. Cultures were incubated at 35°C and 5% CO2 with shaking at 150 rpm for ~48 h.

Following incubation, bacterial cells were pelleted by centrifugation at 3,200 × g for 15 min at 4°C. Pellets were resuspended in 1 mL PBS and stored at -20°C for subsequent DNA extraction. Supernatants were filtered through a 0.22 µm cellulose acetate filter to remove bacterial cells, retaining the phage fraction.

Filtered phage suspensions were used to initiate the next passage by inoculating them at a 1:20 dilution into freshly prepared bacterial cultures as described above. Remaining phage suspensions were stored in glass tubes at 4°C until further use.

This procedure was performed for two sequential passages, resulting in three consecutive culture cycles with ancestral bacterial strains and passaged phages. For the final passage, cultures were scaled up to 35 mL instead of 10 mL to obtain a larger volume of phage suspension for subsequent experiments. Bacterial and phage inoculum ratios were maintained proportionally to ensure identical starting concentrations.

**Synthetic Phage-Bacteria Community Passaging Experiment**

For the experiment including all 12 strains, single-strain pre-cultures were adjusted to an $OD_{600}$ of 0.5 in PBS prior to mixing. For the experiment involving six bacterial communities of different composition (Bicom 1-6, **Fig. 1A**), 50 µL aliquots of single-strain glycerol stocks ($OD_{600}$ = 10) were thawed and adjusted to an $OD_{600}$ of 0.5 in PBS. For all experiments, strains were combined in equal volumes to obtain the desired community composition. An aliquot of each initial community was stored at -80°C for subsequent DNA extraction.

Phage cocktails were prepared by pooling equal volumes of all enriched phage cocktail derived from a given initial phage cocktail (A, B or C) following the phage enrichment experiment.

Initial bacterial communities were inoculated into 7 mL of cfMRS supplemented with the appropriate carbon source (SW or PE) to a final $OD_{600}$ of 0.05. Phage cocktails, or an equivalent volume of cfMRS (control treatment), were added at a 1:20 dilution (350 µL into 7 mL of culture). To monitor growth dynamics, 150 µL aliquots were distributed in triplicate into a 96-well plate for OD measurements. Cultures were incubated at 35°C under 5% $CO_2$ with shaking at 150 rpm for ~48 h.

For passaging, 350 µL of culture (20-fold dilution) was transferred into 7 mL of fresh cfMRS containing the appropriate carbon source. Again, 150 µL aliquots were plated in triplicate for OD monitoring. Cultures were incubated under the same conditions for ~48 h.

After each passage, an aliquot of the culture was stored at -80°C in 40% glycerol. Bacterial cells were pelleted by centrifugation at 3,200 × g for 15 min at 4°C. Supernatants were filtered through 0.22 µm cellulose acetate filters, and the resulting phage suspensions were stored at 4°C in glass tubes. Bacterial pellets were resuspended in 1 mL PBS and stored at -80°C for DNA extraction.

Passaging was repeated for 15 cultures in the 12-strain pilot experiment and for 10 cultures in the experiment with the six Bicoms.

**Plaque Assay**

Plaque assays were performed to assess the lytic activity of phage cocktails obtained throughout the experiments against the 12 *Bifidobacterium* strains used in this study. Assays were conducted using a double-layer agar method consisting of a solid bottom layer and a soft-agar overlay containing bacterial cells.

The bottom layer consisted of MRS agar (1.5% [w/v] agar) supplemented with 2% fructose and 0.05% L-cysteine. The top layer (soft agar) was prepared using MRS broth (BD, #288130) supplemented with 0.55% (w/v) granulated agar, 2% glucose, 2% fructose, 0.1% L-cysteine-HCl, 10 mM $CaCl_2$, and 10 mM $MgCl_2$. The molten soft agar was inoculated with bacterial cells pre-cultured in cfMRS supplemented with SW (final $OD_{600}$ = 0.005) before being poured onto the solidified bottom layer.

After the overlay had solidified, 3-4 µL of phage suspension was spotted onto the surface. When required, phage cocktails were serially diluted in SM buffer using 10-fold dilution steps in a 96-well plate prior to spotting. Once the spots were fully absorbed into the agar, the plates were incubated anaerobically at 35°C for 2-3 days.

**Plaque Titration**

To quantify phage titers longitudinally, plaque assays were performed at the end of each passage using serial 10-fold dilutions of the phage suspension in SM buffer. Dilutions (3 µL) were spotted onto double-layer agar plates prepared as described above, using ESL1060 and ESL1077 as target strains. Plates were incubated anaerobically at 35°C for 2-3 days. The highest dilution yielding visible plaques was recorded and used to calculate phage density (PFU/mL).

**Spot Assays for Community-Level Infectivity Profiling**

To characterize phage infectivity, spot assays were performed against the *Bifidobacterium* strains of this study using a double-layer agar method as described above. Phage suspensions were spotted (3-4 µL) onto plates prepared with each strain and incubated anaerobically at 35°C for 2-3 days. Lytic activity was scored on a qualitative scale from 0 to 3: 0 (no clearing), 1 (weak clearing or individual plaques), 2 (moderate clearing with distinct plaques), and 3 (strong clearing with confluent lysis). This protocol was applied in two contexts. First, to characterize the host range of the enriched phage cocktails, spot assays were performed on all cocktails obtained after the final enrichment passage against each of the 12 strains (Fig. S1E, table S4). Second, to track community-level infectivity profiles longitudinally, spot assays were performed at the end of each passage throughout the experiment with the six Bicoms against the 11 Bifidobacterium strains included in that experiment (Fig. S15, table S16). From these scores, plaque richness (number of strains lysed per passage) and temporal turnover of infectivity profiles were derived.

**Bacterial Growth Dynamics Measurements & Analysis**

Bacterial growth dynamics were assessed by inoculating 150 µL of bacterial culture at $OD_{600}$ = 0.05 into individual wells of a 96-well plate, with each strain or condition performed in triplicate. Plates were incubated at 34°C in a BioTek Synergy H1 microplate reader (Agilent) without shaking. $OD_{600}$ measurements were recorded every 30 min over the entire length of the experiment. Growth curve analysis and visualization were performed using a custom R script. Growth parameters, including maximum growth rate and area under the curve (AUC), were estimated by fitting a logistic growth model to the $OD_{600}$ readings using the R package GrowthCurver v0.3.1 (*47*).

**Sensitivity Analysis for Phage Contamination in Bicom2**

Phage contamination was detected in the no-phage condition of Bicom2 in PE at passages 8-10, evidenced by an unexpected drop in community OD at passage 9 (fig. S11B) and plaque formation at passages 8-10 (fig. S10A). Repeating all analyses after excluding these passages left the effects of phages on alpha, beta, and gamma diversity qualitatively unchanged (fig. S10B), confirming that this contamination did not affect our conclusions.

**Bacterial DNA Extraction**

Following the phage enrichment experiment, a 200 μL aliquot was collected from frozen bacterial suspensions stored in 1 mL PBS. Cells were pelleted by centrifugation at 10,000 × g for 2 min. After centrifugation, 150 μL of supernatant was removed, and the pellet was resuspended in the remaining 50 μL. Then, 150 μL of lysozyme solution (20 mg/mL) was added, and samples were incubated at 37°C for 2 h with shaking at 300 rpm. DNA extraction was then performed using the FastPure Bacteria DNA Isolation Mini Kit (Vazyme) according to the manufacturer's protocol for Gram-positive bacteria. DNA was eluted in 100 μL of 100 mM Tris-HCl.

Following the phage-bacteria passaging experiments, an alternative DNA extraction protocol was employed to efficiently process a large number of samples. A 150 μL aliquot was collected from frozen bacterial suspensions stored in 1 mL PBS. To lyse the cells, 150 μL of G2 buffer and 15 μL of lysozyme (100 mg/mL) were added, and the mixture was incubated at 37°C for 45 min with shaking at 600 rpm. Subsequently, 15 μL of Proteinase K (20 mg/mL) was added, and the samples were incubated at 56°C for 1 h with shaking at 900 rpm. Cell debris was removed by centrifugation at 10,000 × g for 1 min. The supernatant was treated with 4 μL of RNase A (10 mg/mL), and 80 μL of the lysate was transferred to individual wells of a 96-well plate.

To each well, 80 μL of NGS Clean magnetic beads (LabGene, #CNGS0050) were added and mixed by pipetting. The plate was incubated at room temperature for 10 min, then placed on a magnetic rack. Once the beads had collected on the side of the wells, the supernatant was discarded, and the beads were washed twice with 180 μL of 80% ethanol. After the ethanol had completely evaporated, DNA was eluted in 50 μL of 10 mM Tris-HCl and transferred to new tubes following a 5 min incubation with the beads.

For all protocols, extraction blanks consisting of PBS buffer were included in each DNA extraction batch. DNA concentrations were quantified using a Qubit fluorometer, and samples were stored at −20 °C or −80 °C until further use.

**Viral DNA Extraction**

We extracted viral DNA from the initial and enriched phage cocktails to determine their community composition. We additionally sequenced phages from Passages 1, 3, 4, 5, 6, 9, 12, and 15 of the 12-strain passaging experiment. Although the sequencing data of that experiment was not included in any community-level analysis, it was used to assemble viral genomes and improve the clustering of viral sequences into vOTUs (see *Methods, Shotgun Metagenomics data processing*).

To extract viral DNA, 750 μL of phage suspension was incubated with 27 μL of DNase I reaction buffer (New England Biolabs, #B0303S), 10 μL of DNase I (10 mg/mL), and 10 μL of RNase A (10 mg/mL) at 37°C for 15 min to degrade free bacterial DNA and RNA.

The reaction was stopped by adding 150 µL of EDTA buffer (133.3 µL 0.5 M EDTA, 8.3 µL 1 M Tris-HCl pH 7.5, 8.3 µL $ddH_2O$) and incubating at 65°C for 10 min. Viral capsids were then lysed by adding 75 µL of 10% SDS and 12.5 µL of Proteinase K (20 mg/mL), followed by incubation at 55°C with shaking at 300 rpm for 1 h. DNA was purified by adding 1 mL of phenol:chloroform:isoamyl alcohol (25:24:1, stabilized, saturated with 100 mM Tris-EDTA, pH 8.0; Fisher Scientific, #10535691), mixing vigorously, and centrifuging at 16,000 × g for 10 min. The aqueous phase was transferred to a new tube, and the phenol:chloroform:isoamyl alcohol extraction was repeated once, followed by a chloroform extraction. DNA was precipitated by adding 550 µL of ice-cold isopropanol, 55 µL of 3 M sodium acetate (pH 5.2), and 1 µL of GenElute Linear Polyacrylamide (Sigma, #56575-1ML). Samples were incubated at -20°C for 1 h and centrifuged at maximum speed for 30 min at 4°C. The pellet was washed with ice-cold 80% ethanol, centrifuged again for 15 min at 4°C, and resuspended in 30 µL of 10 mM Tris-HCl. For all protocols, extraction blanks consisting of SM buffer were included in each DNA extraction batch. DNA concentration was quantified using a Qubit fluorometer, and samples were stored at -20°C until further use.

**16S rRNA Gene Amplicon Sequencing of Bacterial Communities**

To characterize bacterial community composition, we performed full-length 16S rRNA gene amplicon sequencing. For the 249 samples of the passaging of the six Bicoms (240 cultures, 6 inoculums and 3 extraction blanks), multiplexed full-length 16S rRNA gene amplicon libraries were prepared following the *“Preparing Kinnex™ libraries from 16S rRNA amplicons”* protocol (Version 06, Pacific Biosciences). PCR amplifications were carried out using KAPA HiFi HotStart ReadyMix (KAPA Biosystems) and a pre-arrayed 96-well barcoded primer plate. Each well contained a unique combination of forward (5′-CTACACGACGCTCTTCCGATCT/barcode/AGRGTTYGATYMTGGCTCAG-3′) and reverse (5′-AAGCAGTGGTATCAACGCAGAG/barcode/RGYTACCTTGTTACGACTT-3′) primers, generating 96 distinct barcode pairs according to the manufacturer’s recommendations. Amplicon concentrations were not normalized prior to sequencing. To accommodate the 365 samples of the 12-strain passaging experiment (360 cultures, 1 inoculum, and 4 extraction blanks), 16S rRNA gene amplicon libraries and PCR amplification were performed as described above. Equimolar amounts of individual amplicons were pooled and subjected to the Kinnex™ concatenation protocol, followed by SMRTbell® library preparation using the SMRTbell Prep Kit 3.0 to obtain concatenated and multiplexed full-length 16S rRNA gene amplicon libraries. For all protocols, library preparation and sequencing were performed on a PacBio Sequel II system at the Genomic Technologies Facility of the University of Lausanne.

**Shotgun metagenomics sequencing**

Viral community composition was assessed using shotgun metagenomic sequencing. Metagenomic libraries were prepared using Nextera DNA Flex Library Prep Kit

(Illumina) with unique dual indices (UDIs) and sequenced on an AVITI platform (Element Biosciences) to generate 150 bp paired-end reads. Library preparation and sequencing were performed at the Genomic Technologies Facility of the University of Lausanne.

**Bacterial phylogeny**

*Bifidobacterium* genomes (table S1) were annotated using DRAM v1.4.0 (*48*). Orthogroups were inferred from the proteomes using OrthoFinder v3.0.1 (*49*) with the "-m MSA" option to generate multiple sequence alignments of 991 single-copy orthogroups conserved across genomes. The resulting concatenated alignment was trimmed with trimAl v1.5.0 (*50*). A maximum-likelihood phylogeny was then reconstructed using IQ-TREE v3.0.1 (substitution model: LG+F+I+G4; bootstraps: 100; (*51*). The final phylogenetic tree was visualized in R.

**16S rRNA Gene Amplicon sequence analysis**

Amplicon sequencing raw reads were processed using DADA2 v1.26.0 (*52*) to infer amplicon sequence variants (ASVs) and their read counts across samples. Briefly, primers were removed from raw reads using the *removePrimers* function. Reads were then filtered to retain sequences between 1,400 and 1,600 bp. Finally, reads were dereplicated and denoised using the binnedQualErrfun error model. The complete analysis workflow is available at: https://github.com/MalickNdiye/PHIFIDO.

A custom 16S rRNA reference database was constructed by combining the SILVA v138.2 16S rRNA database with 16S rRNA sequences extracted from the genomes of the 12 *Bifidobacterium* strains used in this study using barrnap v0.9. Inferred ASVs were matched against this database to identify strain-specific 16S rRNA gene variants (table S18). These assignments were used to infer strain-level counts based on the ASV count table generated by DADA2.

Strain relative abundance was calculated as the proportion of reads assigned to each strain relative to the total reads per sample. ASVs with <10% prevalence and < 10% maximum relative abundance across samples within the same sequencing run were removed (fig. S18A-B), as these likely reflect contaminations, sequencing artifacts, or residual chimeras. In the 12-strain passaging experiment, additional ASVs assigned to the genus *Bifidobacterium* but not to any of the included strains were removed manually, as they likely represented sequencing errors (fig. S18A). In the passaging experiment with the six Bicoms, ASVs belonging to strains not included in a given community were sporadically detected at low average relative abundances (<1%) across samples (table S19). Several of these ASVs were found also in the DNA extraction blanks (table S10). The relative abundance of these samples was strongly negatively correlated with library size (Spearman's rho < -0.5; Fig. S18C). This means that with fewer total reads, more of these unexpected ASVs were detected, indicating contamination likely occurred during DNA extraction or library preparation, rather than during the experiment. Thus, they were removed manually. Absolute strain abundance was estimated by multiplying relative abundance by the Area Under the growth Curve as inferred by OD readings.

**Shotgun Metagenomics Data Processing**

Raw metagenomic reads were trimmed and filtered to remove low-quality bases and short sequences using Trimmomatic v0.36 (*53*) with the following parameters: *PE ILLUMINACLIP:NexteraPE-PE.fa:2:30:10 LEADING:25 TRAILING:25 MINLEN:40*. Quality-filtered reads were assembled using MEGAHIT v1.2.9 .

Viral contigs were identified from metagenomic assemblies using VIBRANT v1.2.1 (*54*). Additionally, prophages were detected in the genomes of the 12 *Bifidobacterium* strains using geNomad v1.8.1 (*55*). All viral contigs (from both viromes and prophage predictions) were quality assessed using CheckV v1.0.1 (56). Only contigs ≥5 kb classified as “Medium-quality” or higher were retained for downstream analyses. Phage lifestyle (temperate or virulent) was predicted using BACPHLIP v0.9.6, and taxonomy was assigned using PhaBOX2 v2.1.9 (--task phagcn).

Following MIUViG guidelines (Roux et al., 2019 (*57*)), retained viral contigs were dereplicated into viral operational taxonomic units (vOTUs) at 95% average nucleotide identity (ANI) and 85% alignment fraction (AF) as in Ndiaye et al. (2025) (*35*), using dRep v3.4.0 (*58*) (parameters: *--S_algorithm ANImf -pa 0.5 -sa 0.95 -nc 0.85 -l 5000 -N50W 0 -sizeW 1 --clusterAlg average --ignoreGenomeQuality*). For each vOTU, the longest contig with the highest CheckV quality score was selected as the representative genome.

Quality-filtered reads were mapped against a combined database of representative vOTU sequences using Bowtie2 v2.5.1 (*59*) in sensitive mode. Community profiling was performed using inStrain v1.10 (*60*). Only vOTUs with coverage breadth ≥0.7 were considered as present in a given community.

**In-Silico Detection of Prophage Induction**

To detect prophage induction, we mapped sequencing reads from each sample to the genomes of the 12 Bifidobacterium strains used in this study and calculated per-position coverage profiles using the bbsplit function from bbmap v39.13. Prophage coordinates were obtained from the geNomad results. For each contig in each sample, we computed the mean genome-wide coverage as a baseline. A prophage was considered induced if its median coverage exceeded 1.5 times the genome-wide average. This analysis was restricted to strains for which phages were recovered in the control samples during phage enrichment (ESL0819, ESL0198, ESL0170, mixSW, mixPE).

**Phage-Host Prediction via CRISPR Spacers-to-Protospacers Match**

To assign hosts to the vOTUs recovered in this study, we used CRISPR spacer-to-protospacer matches. CRISPR spacers were sourced from a database of honeybee bacterial spacers established in Ndiaye et al. (2025) (*35*) and mapped against the vOTU sequences using BLASTn with the "blastn-short" task. A spacer-to-protospacer match was accepted only when the alignment spanned the full length of the spacer query with no more than 2 mismatches.

**Statistical Analysis**

Alpha-diversity over time: For bacterial communities, alpha-diversity was quantified as the Hill number of order 1 ($H_1$ = exp(H'), where H' is the Shannon entropy), which expresses diversity in units of effective species number. For phage communities, the breadth of the infectivity profile was quantified as plaque richness, defined as the number of bacterial strains against which plaques were detected at a given passage in the spot assay.

$H_1$ and plaque richness over time were modeled using Generalized Additive Models (GAMs) fitted by REML using the mgcv v1.9-3 package in R. For each response variable, the error distribution family was selected by inspecting the empirical distribution of the response variable and the relationship between group-level means and variances. Non-linear temporal dynamics were captured using penalised regression splines over passage number, and the random effect structure was chosen to account for the repeated-measures over time and paired design of the experiment (same communities in different conditions). For each analysis, a candidate set of models was evaluated, varying in the complexity of the temporal smooth structure, the random effects specification, and the inclusion of treatment terms. A null model excluding all predictors was always included as null hypothesis. Full details of the candidate models tested, including their formula, family, and random effect structure, are provided in table S20-22.

Model selection was performed using a two-step procedure. First, candidate models were ranked by AIC, then residual diagnostics were assessed for each model using DHARMa v0.4.7 R package, testing for uniformity, dispersion, outliers, and temporal autocorrelation. BIC was used as a secondary parsimony criterion. The selected model was the lowest-AIC model whose residuals passed all diagnostic checks. Full diagnostic scores for all candidate models are provided in table S23-25.

Estimated marginal means and pairwise contrasts were calculated from the selected models using the emmeans v2.0.2 R package. All p-values were corrected for multiple comparisons using the FDR method. Effect sizes were reported as Glass's Δ, calculated as the estimated marginal difference divided by the standard deviation of the relevant reference group (always SW and/or NO-PHAGES).

Beta-diversity: To characterize compositional turnover over time, we calculated pairwise Bray-Curtis dissimilarities between all combinations of timepoints within each community, resource condition, and phage treatment, using relative strain abundances and the vegdist function from the vegan v2.7-3 package. For each community, we fitted a linear model regressing Bray-Curtis dissimilarity against the temporal distance between passages (Δtime). From this model, we extracted two summary statistics: the dissimilarity at Δtime = 1 (intercept + slope, representing short-term passage-to-passage turnover) and the slope (representing the rate of community divergence over time).

Both statistics were modeled using linear mixed-effects models (lme4 v2.0-1) with media condition, phage treatment, and their interaction as fixed effects, and community identity as a random intercept. The same approach was used for the phage infectivity profiles, using resource condition as the only fixed effect. Estimated marginal means, pairwise contrasts, and Glass's Δ effect sizes were computed as described for the alpha-diversity analysis, with FDR correction applied to all p-values.

Oscillations: To quantify strain-level temporal instability, we developed an oscillation score (CV_RMSE) measuring the deviation of each strain's abundance trajectory from non-oscillating growth. For each strain in each community and condition, we fitted three candidate models to the longitudinal absolute abundance data (relative abundance x community AUC): a logistic growth model (SSlogis), an asymptotic model (SSasymp), and a linear model as fallback. The best-fitting model was selected sequentially, falling back to simpler models when convergence failed. The oscillation score was then calculated as the root-mean-square error of the residuals divided by the mean abundance (CV_RMSE), such that strains reaching carrying capacity, going extinct, or remaining at steady state yield values near zero, while strains exhibiting oscillatory dynamics yield higher values.
Oscillation scores were averaged per strain across communities and modelled using a linear mixed-effects model with resource condition, phage treatment, and their interaction as fixed effects, and strain identity as a random intercept.
Estimated marginal means, pairwise contrasts, and Glass's Δ effect sizes were computed as described for the alpha-diversity analysis, with FDR correction applied to all p-values.

Gamma-diversity: To estimate the effective species pool maintained across the experiment, we calculated temporal gamma diversity for each community as the $H_1$ computed on mean relative abundances across all passages. This integrates both the richness and the average relative contributions of strains over time, providing a single estimate of the effective number of species persisting throughout the experiment.
Temporal gamma diversity was modelled using a linear mixed-effects model (lme4 v2.0-1) with resource condition, phage treatment, and their interaction as fixed effects, and community identity as a random intercept.
Estimated marginal means, pairwise contrasts, and Glass's Δ effect sizes were computed as described for the alpha-diversity analysis, with FDR correction applied to all p-values.

**Mathematical Model**

This section gives a shortened account of the model; a complete description is available in the Supplementary Material *Model Description*. We built a mathematical model describing changes in abundance of bacterial species, phages, and nutrients under resource competition and phage predation. The model includes $Ns$ species, $Np$ phages, and $Nr$ resources. Species $i$ grows on resource α at rate $\mu_{i\alpha}$. Phage $k$ infects species *i* at per-capita adsorption rate $\eta_{ki}$ (drawn from a Gamma distribution), with burst rate $\beta$. This yields a phage-induced death rate of $\beta * \eta_{ki}$ per bacterial species. Full dynamical equations are in the extended methods.
For simplicity, we consider a chemostat setup with a constant dilution rate $D$ = 0.1 $h^{-1}$ instead of a serial transfer protocol. Resources are supplied at 5 mM and depleted by

dilution and consumption. Each species is characterized by its maximum growth rate and total susceptibility to all phages.

To represent sucrose water and pollen extract, we make two empirically motivated choices. Sucrose water has fewer usable resources (smaller $Nr$) and a denser growth-rate matrix. Instead, pollen extract has broader niche breadth and a sparser matrix, with identical maximal growth rates. Matrix density is set by the shape parameter of the Gamma distribution from which growth rates are drawn (sucrose water: 2; pollen extract: 0.25). Simulations use $Np$ = 24, $Ns$ = 12, and $Nr$ = 6 (sucrose water) or 30 (for pollen extract). The numbers of phages and bacterial species were considered alongside the experimental numbers, and the choice of $Nr$ does not affect the qualitative findings.

### Model description: Niche Overlap Determines Phage Predation Impact on Bacterial Diversity

Malick Ndiaye*, Thania Sbaghdi*, Lucie Kesner, Prajwal Padmanabha, Meline Garcia, Ena Bochaton, Sara Mitri, Philipp Engel

#### Model definition

We built a consumer-resource model simulating the dynamics of bacterial species, phages, and nutrient concentrations in a chemostat. The model considers $N_s = 12$ bacterial species, $N_p = 24$ phages, and $N_r$ resources (where $N_r$ depends on the resource environment; see below). Bacterial abundances are denoted $n_i$, resource concentrations $r_\alpha$, and phage abundances $p_k$. The dynamics are governed by

$$\frac{dn_i}{dt} = n_i \left( \sum_\alpha \mu_{i\alpha} r_\alpha - \sum_k \eta_{ki} p_k - D \right) \tag{1}$$

$$\frac{dr_\alpha}{dt} = D(S_\alpha - r_\alpha) - r_\alpha \sum_i \mu_{i\alpha} n_i \tag{2}$$

$$\frac{dp_k}{dt} = p_k \left( \beta \sum_i \eta_{ki} n_i - D \right) \tag{3}$$

where $\mu_{i\alpha}$ is the uptake rate of species $i$ on resource $\alpha$, $\eta_{ki}$ is the adsorption rate of phage $k$ on species $i$, $D$ is the dilution rate, $S_\alpha$ is the supply concentration of resource $\alpha$, and $\beta$ is the burst size (number of phage progeny produced per lysis event). Bacterial growth on each resource is linear in resource concentration (non-saturating uptake).

#### Parameter values

Each bacterial species $i$ is characterized by a maximum growth rate $g_i$ and a total phage susceptibility $\sigma_i$. These are drawn jointly from a bivariate log-normal distribution, where the covariance matrix has a standard deviation of 0.5 in each marginal and zero off-diagonal correlation. This means that the maximum growth rate and total phage susceptibility are drawn independently. This independence reflects no external imposition of a growth-defense tradeoff. So, any relationship between competitive ability and vulnerability to phages arises solely from the ecological dynamics.

The per-resource uptake rates $\mu_{i\alpha}$ are constructed by drawing raw entries from a Gamma distribution, $\mu_{i\alpha}^{\text{raw}} \sim \text{Gamma}(k, \theta)$, where $k$ is the shape parameter. Each row is then normalized so that the total growth rate of species $i$ across all resources equals its assigned maximum growth rate:

$$\mu_{i\alpha} = \mu_{i\alpha}^{\text{raw}} \times \frac{g_i}{\sum_\alpha \mu_{i\alpha}^{\text{raw}}} \tag{4}$$

The shape parameter $k$ controls the sparsity of the uptake matrix and therefore the degree of metabolic niche overlap among species. A high shape value ($k = 2$, used for sucrose water) produces dense, relatively uniform uptake vectors where all species consume all resources at comparable rates, resulting in high niche overlap. A low shape value ($k = 0.25$, used for pollen extract) produces sparse, skewed uptake vectors where each species specializes on a small subset of resources, resulting in low niche overlap.

For a given community, the total growth rate of each species ($\sum_\alpha \mu_{i\alpha} = g_i$) is constant across resource environments. What changes between environments is how this total is distributed across resources.

The adsorption rates $\eta_{ki}$ are constructed similarly. Raw entries are drawn from $\eta_{ki}^{\text{raw}} \sim \text{Gamma}(0.5, 2.5)$. Each column (corresponding to a species $i$) is then normalized so that the total rate across all phages equals the species assigned total susceptibility:

$$\eta_{ki} = \eta_{ki}^{\text{raw}} \times \frac{\sigma_i}{\sum_k \eta_{ki}^{\text{raw}}} \tag{5}$$

The shape parameter 0.5 produces a moderately sparse infection matrix, reflecting heterogeneity in phage host range. The infection matrix is the same for a given community regardless of the resource environment.

#### Resource environments

Sucrose water and pollen extract differ in two respects: the number of resources supplied and the structure of the growth rate matrix.

**Sucrose water** $N_r^{\text{SW}} = 6$ resources are supplied (the 6 resources with the highest total consumption $\sum_i \mu_{i\alpha}$ across species), each at concentration $S_\alpha = 25$ ($= S_{\text{base}} \times N_r / N_r^{\text{SW}}$, where $S_{\text{base}} = 5$). All remaining resources are effectively absent ($S_\alpha = 10^{-30}$). The growth rate matrix has shape parameter $k = 2$ (dense, high niche overlap).

**Pollen extract** All $N_r = 30$ resources are supplied uniformly at concentration $S_\alpha = 5$. The growth rate matrix has shape parameter $k = 0.25$ (sparse, low niche overlap).

This construction ensures that the total resource supply ($\sum_\alpha S_\alpha = 150$) is equal across environments. The environments differ in resource diversity and metabolic overlap, not in total resource availability.

For a given growth rate matrix, we quantify metabolic niche overlap as the mean pairwise cosine similarity of species' uptake vectors:

$$\rho = \frac{1}{N_s(N_s - 1)} \sum_{i \neq j} \frac{\sum_\alpha \mu_{i\alpha} \mu_{j\alpha}}{\sqrt{\sum_\alpha \mu_{i\alpha}^2 \sum_\alpha \mu_{j\alpha}^2}} \tag{6}$$

This measure ranges from 0 (completely orthogonal uptake profiles) to 1 (identical uptake profiles). Higher niche overlap implies stronger exploitative competition.

#### Simulation details

To maintain positivity of all variables during integration, we perform a log-transformation, defining $x = \log(n_i)$, $y = \log(r_\alpha)$, $z = \log(p_k)$, and integrating the transformed system. Any variable whose log-abundance drops below $-35$ (corresponding to an abundance of approximately $6 \times 10^{-16}$) is considered extinct and its derivative is set to zero for the remainder of the simulation. This threshold is sufficiently low that any species or phage reaching it is effectively lost from the community.

The system is integrated for a total time of 1000 arbitrary units using 100 evenly spaced output timepoints. We use the implicit Radau method (in scipy.integrate.solve_ivp) with relative tolerance $10^{-4}$ and absolute tolerance $10^{-9}$, chosen to handle the stiff dynamics that arise from the phage-bacteria interaction.

Initial conditions are: $n_i(0) = 1$ for all species, $r_\alpha(0) = S_\alpha$ (resources at supply concentration), $p_k(0) = 0.1$ for all phages in the phage-present condition, and $p_k(0) = 10^{-30}$ (effectively absent) in the phage-absent condition.

For each parameter combination, 100 random communities are simulated (50 for the niche overlap sweep). Each community consists of a fresh random draw of maximum growth rates and total susceptibilities, a growth rate matrix $\mu$ for each environment, and a shared infection matrix $\eta$. All four conditions of the $2 \times 2$ factorial design (sucrose water/pollen extract $\times$ phage/no phage) are simulated for each community, ensuring that differences between conditions are attributable to the treatment rather than to variation across community draws.

The phage effect on diversity is quantified as $\Delta_{\text{Hill}} = H_{\text{phage}} - H_{\text{no phage}}$, computed separately for each community and environment.

#### Diversity measures

Diversity is computed from the last 10% of the simulation trajectory (last 10 timepoints) to exclude transient dynamics. At each timepoint in this window, we compute the relative abundances $f_i = n_i / \sum_j n_j$ and the Hill diversity (exponential of Shannon entropy):

$$H = \exp\left(-\sum_i f_i \log f_i\right) \tag{7}$$

Species with abundance below $10^{-2}$ at a given timepoint are considered below the survival threshold and included in the calculation at their actual (low) abundance. The reported diversity is the mean Hill diversity across all timepoints in the last 10% window ($\alpha$-diversity).

The oscillation score quantifies temporal variability in species abundances. For each surviving species (endpoint abundance $> 10^{-2}$), we fit a second-degree polynomial to its abundance trajectory over the last 10% of the simulation to remove any residual approach to equilibrium. The oscillation score for that species is the standard deviation of the residuals (observed minus fitted) divided by the mean abundance over the window. The reported oscillation score is the mean across all surviving species. A score near zero indicates steady-state dynamics; larger values indicate sustained oscillations or irregular fluctuations.

#### Robustness analyses

To assess whether the main model findings depend on the specific parameter values chosen, we varied the model parameters across 24 parameter combinations. Specifically, we tested:

- Is sucrose water having more diversity gain due to lower number of supplied resources or higher niche overlap: we gave pollen extract the same number of resources as sucrose water (6 instead of 30) while retaining its low niche overlap.
- Is no trade-off in growth-susceptibility driving any of the diversity or oscillation results: we imposed a positive or negative correlation between growth rate and phage susceptibility ($\rho_{g\sigma} = -0.5$ and 0.9; default $\rho_{g\sigma} = 0$, Table 1).
- Do phage burst size or species growth/infectivity traits make both media show same effects: we varied burst size from $\beta = 5$ to $\beta = 100$, and the mean log growth rate and mean log susceptibility across nine combinations ($\mu_g \in \{-2, -1, 0\}$, $\mu_\sigma \in \{-4, -3, -2\}$).
- Does the differing carbon ratio between pollen and sucrose water impact results: we varied the pollen-to-sugar total carbon ratio from 0.5 to 2.0 (default 1.0).

Across all 24 parameter configurations, the three main model results were robust (Fig. S17). The ordering $\Delta^{\text{SW}}_{\text{Hill}} > \Delta^{\text{PE}}_{\text{Hill}}$ all configurations except the ones where pollen extract had

Table 1: Summary of model parameters.

| # | Parameter | Symbol | Value |
|---|---|---|---|
| 1 | Number of species | $N_s$ | 12 |
| 2 | Number of phages | $N_p$ | 24 |
| 3 | Number of resources | $N_r$ | 30 (6 supplied in SW, 30 in PE) |
| 4 | Dilution rate | $D$ | 0.1 |
| 5 | Burst size | $\beta$ | 10 |
| 6 | Base supply concentration | $S_{\text{base}}$ | 5 |
| 7 | Mean log max growth rate | $\mu_g$ | $-1$ |
| 8 | Mean log max susceptibility | $\mu_\sigma$ | $-3$ |
| 9 | Std. dev. of log traits | $\sigma_{\text{trait}}$ | 0.5 |
| 10 | Growth–susceptibility correlation | $\rho_{g\sigma}$ | 0 |
| 11 | $\mu$ shape, sucrose water | $k_{\text{SW}}$ | 2 |
| 12 | $\mu$ shape, pollen extract | $k_{\text{PE}}$ | 0.25 |
| 13 | $\eta$ shape | $k_\eta$ | 0.5 |
| 14 | Gamma scale (all matrices) | $\theta$ | 2.5 |
| 15 | Simulation time | $T$ | 1000 a.u. |
| 16 | Output timepoints | — | 100 |
| 17 | Extinction threshold (log) | — | $-35$ |
| 18 | Survival threshold | — | $10^{-2}$ |
| 19 | Communities per condition | — | 100 (50 for sweeps) |

high niche-overlap (Fig. S17A). Phages induced stronger oscillations in sucrose water than in pollen extract in 84% of all pooled communities (Wilcoxon $p < 0.001$; Fig. S17B), with reversals concentrated in configurations with very high burst size. The positive cross-media correlation between $\Delta_{\text{Hill}}^{\text{SW}}$ and $\Delta_{\text{Hill}}^{\text{PE}}$ was significant in all 24 configurations (median Pearson $r = 0.35$, all $p < 0.05$; Fig. S17C).

Niche overlap was the only parameter whose manipulation changed the qualitative pattern of phage effects: equalizing niche overlap across environments eliminated the media-dependent difference (main text Fig. 5), while no other parameter change did so.

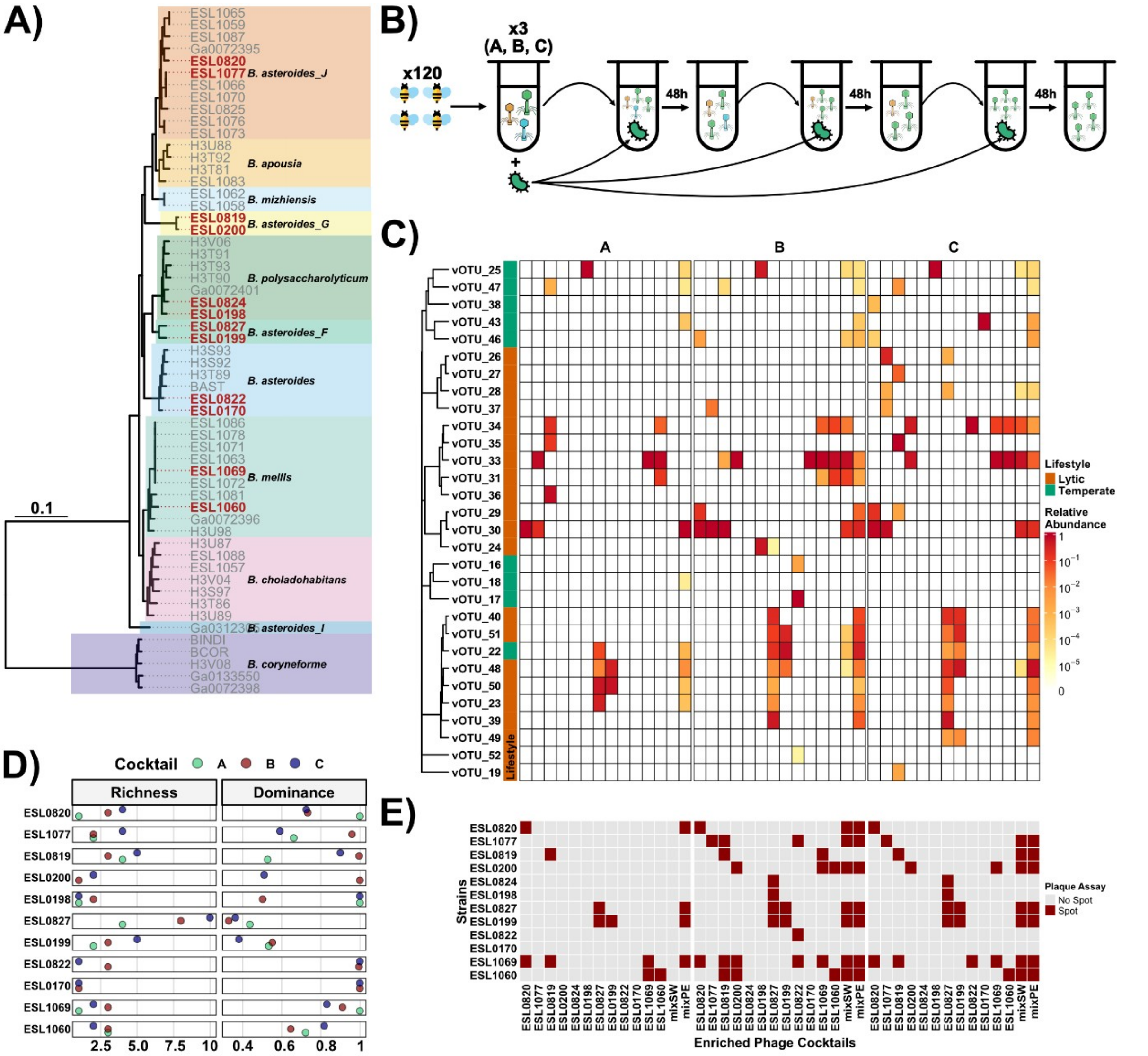


**Fig. S1: Assembly of a diverse synthetic *Bifidobacterium*-phage community.**

(**A**) Maximum-likelihood phylogenetic tree of *Bifidobacterium* strains from the honeybee gut based on multiple sequence alignment of 991 single-copy orthogroups. The 12 strains selected for the experiments in this study are highlighted in red. Colored backgrounds denote *Bifidobacterium* species. Scale bar represents 0.1 substitutions per site. Tree root is not shown. (**B**) Phage enrichment workflow. Three phage cocktails (A, B, C) were independently obtained from gut homogenates pooled from 120-170 honeybees each and enriched by successive 48h co-incubations with each of the 12 individual *Bifidobacterium* strains or with a mix of all 12 strains in sucrose water (SW) or pollen extract (PE). Enrichment was performed over three successive passages, each lasting 48h. (**C**) Heatmap of vOTU relative abundances across enriched phage cocktails. Rows correspond to the 21 enriched vOTUs identified by shotgun metagenomic sequencing of each enriched phage cocktail. Columns correspond to individual enrichment cultures, grouped by cocktail of origin and by the bacterial strain or strain mix used for enrichment, using the same order as in (E). Color indicates relative abundance on a log scale. The sidebar indicates predicted phage lifestyle. (**D**) Richness (number of vOTUs detected) and dominance (Simpson's dominance index) of enriched cocktails for each bacterial strain, colored by cocktail of origin. (**E**) Plaque assay matrix showing lytic activity of all enriched cocktails (columns) against all 12 bacterial strains (rows).

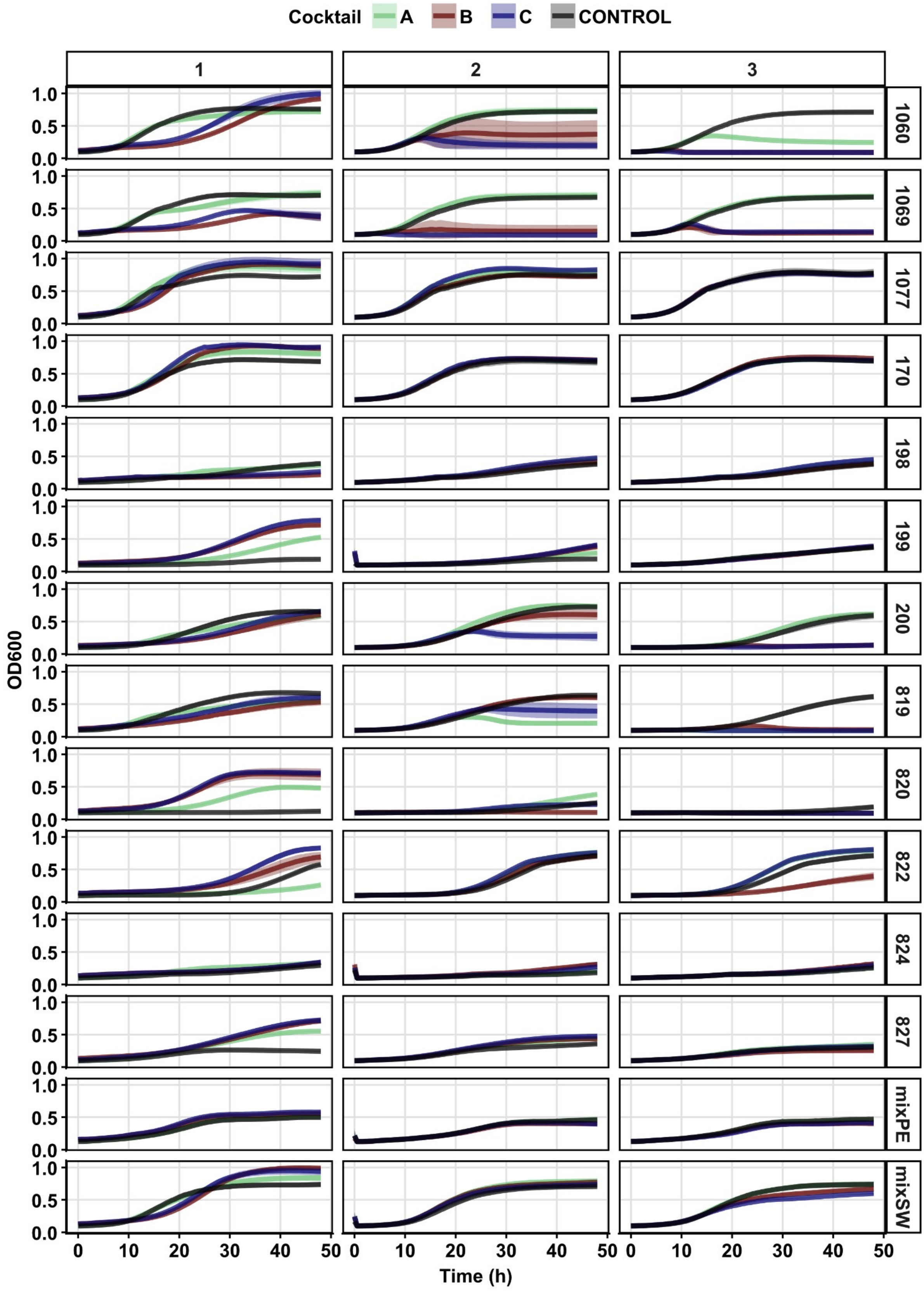


**Fig. S2: Growth curves of individual *Bifidobacterium* strains and strain mixes during phage enrichment.**

OD600 measured over 48h for each bacterial strain or strain mix (rows) across three successive enrichment passages (columns 1-3), in the presence of phage cocktails A (green),

B (red), or C (blue), or without phages (CONTROL, black). Shaded ribbons show the standard deviation across technical replicates (n=3).

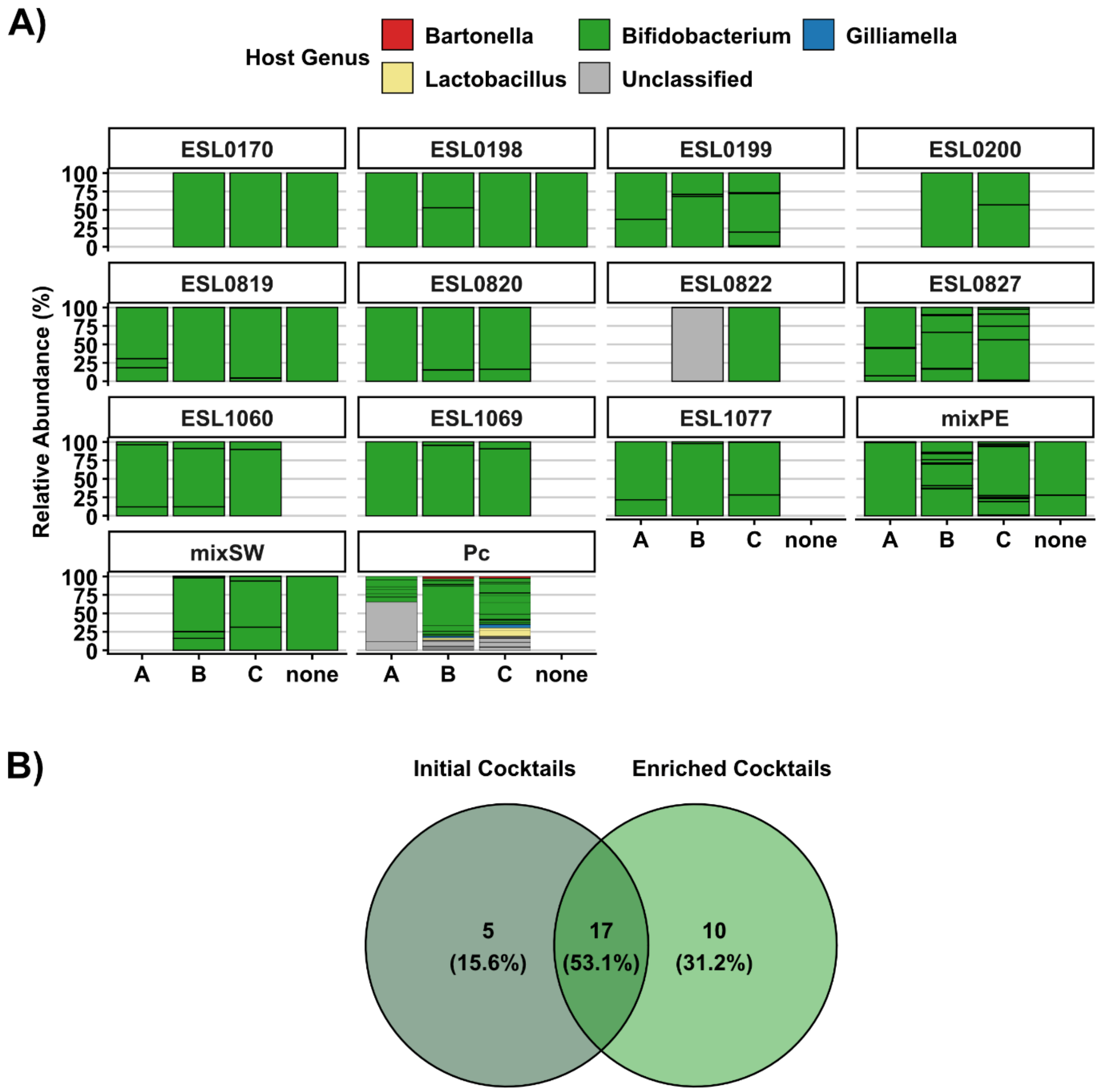


**Fig. S3: Enriched phage cocktails are dominated by Bifidobacterium-targeting vOTUs.** (**A**) Composition of enriched phage cocktails based on predicted host genus. Each panel corresponds to a bacterial strain or strain mix used for enrichment. Bars show the relative abundance of vOTUs in each of these enrichments across the three different cocktails of origin (A, B, C) and the no-phage control (none). PC, initial phage cocktail; mixSW, 12-strain mix in sucrose water; mixPE, 12-strain mix in pollen extract. (**B**) Venn diagram showing the overlap between *Bifidobacterium-targeting* vOTUs detected in the initial (non-enriched) phage cocktails and vOTUs recovered after enrichment.

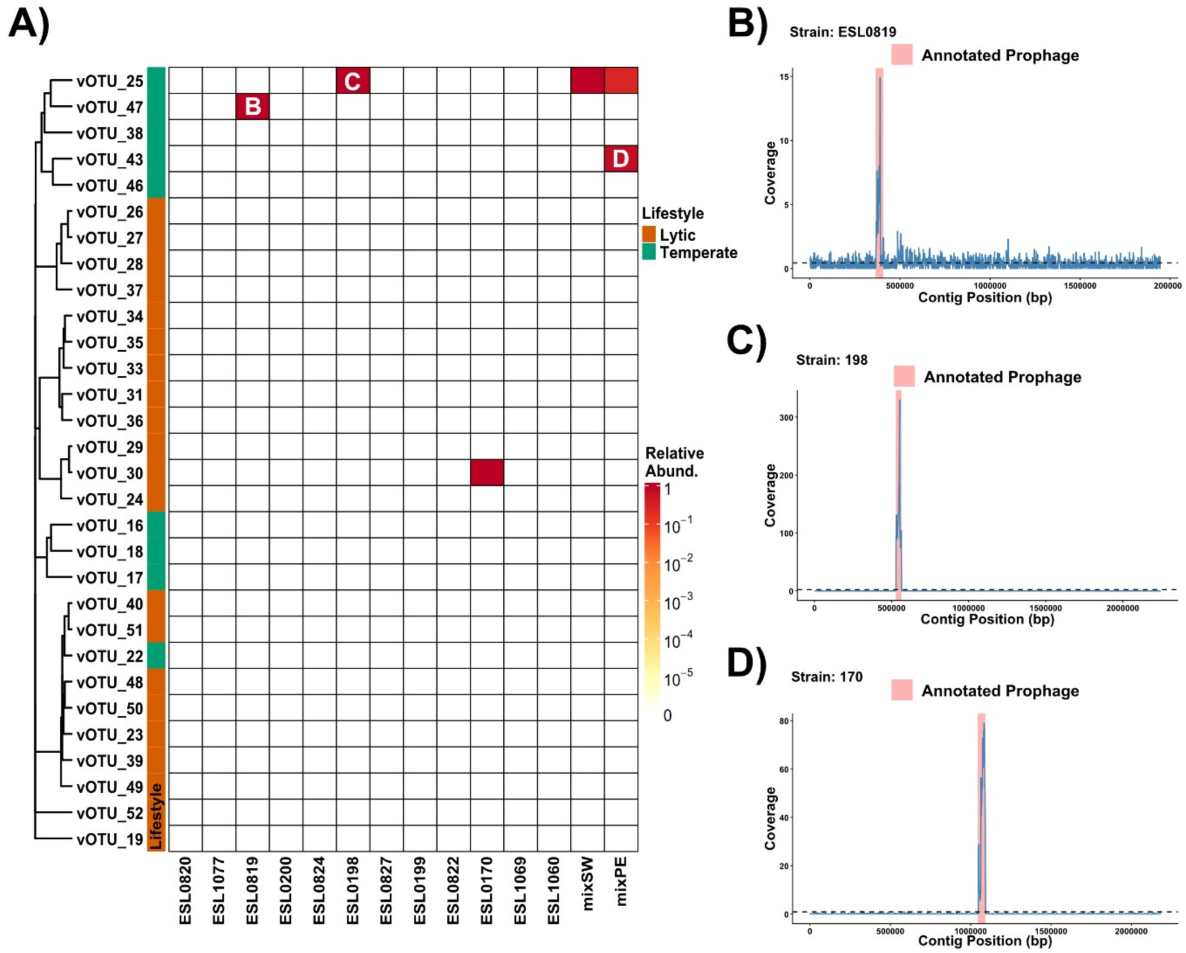


**Fig. S4: vOTUs detected in no-phage controls during phage enrichment.**

**(A)** Heatmap of vOTU relative abundances across enriched phage cocktails, as in fig. S1C. Panels (B-D) indicate the three prophages detected in no-phage control samples. The sidebar indicates predicted phage lifestyle. **(B-D)** Genomic coverage profiles supporting prophage induction in strains (C) ESL0819, (D) ESL0198, and (E) ESL0170. The blue line shows sequencing coverage (1 kb rolling mean) along the entire bacterial contig. The pink shaded region indicates the annotated prophage coordinates. The dashed line shows the genome-wide average coverage.

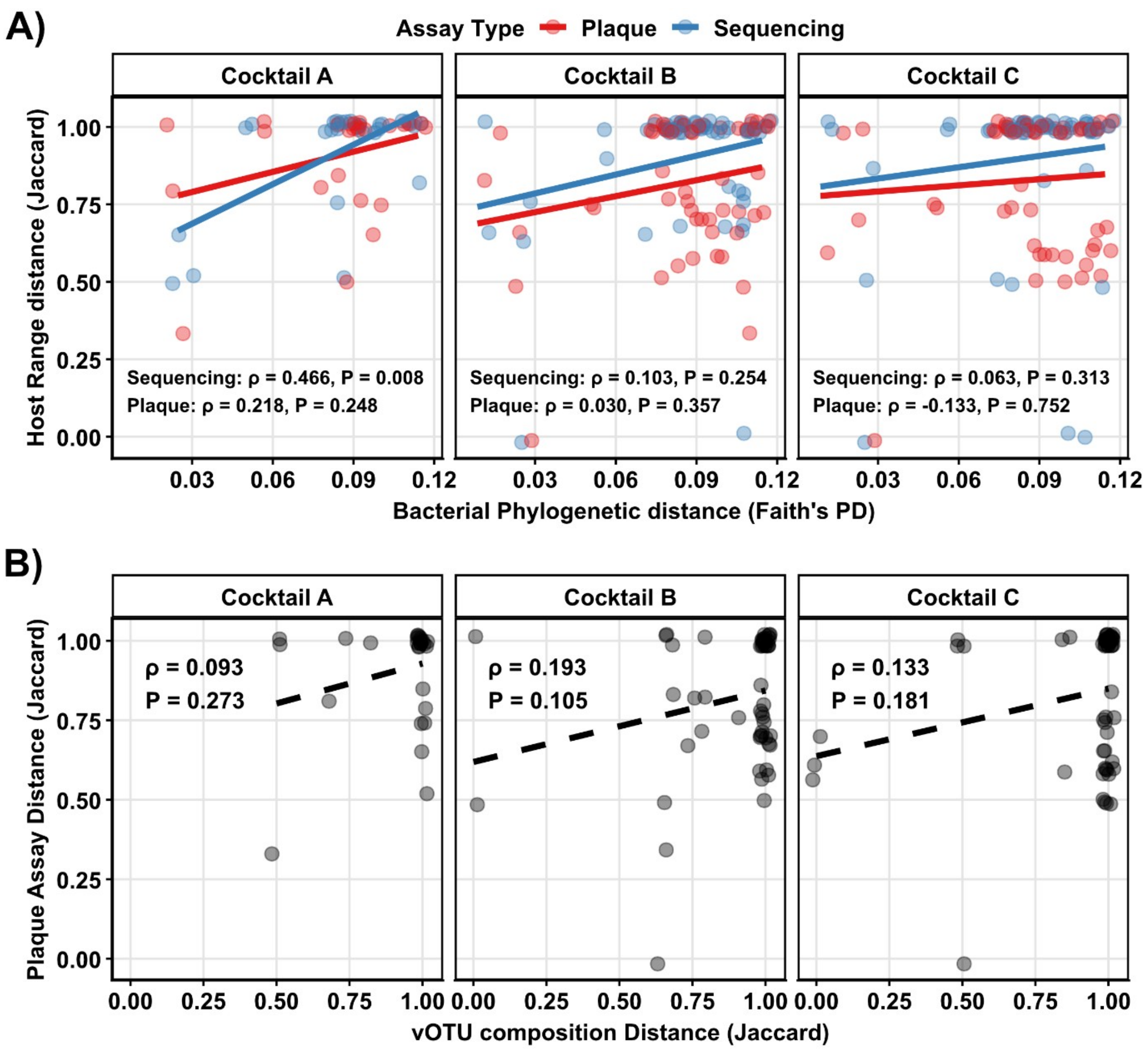


**Fig. S5**: **Phage host range is not predicted by bacterial phylogenetic distance or vOTU composition.**

(**A**) Correlation between bacterial phylogenetic distance (Faith's PD, x-axis) and phage host range distance (Jaccard, y-axis) for each pair of bacterial strains, shown separately for each cocktail of origin (A, B, C). Host range distance was measured either by plaque assay (red) or by vOTU composition from metagenomic sequencing (blue). Lines show linear regression fits. Spearman correlation coefficients (ρ) and p-values are indicated for each measure. (**B**) Correlation between vOTU composition distance (Jaccard, x-axis) and plaque assay distance (Jaccard, y-axis) for each pair of enriched cocktails, shown separately for each cocktail of origin (A, B, C). Dashed lines show linear regression fits. Spearman correlation coefficients and p-values are indicated.

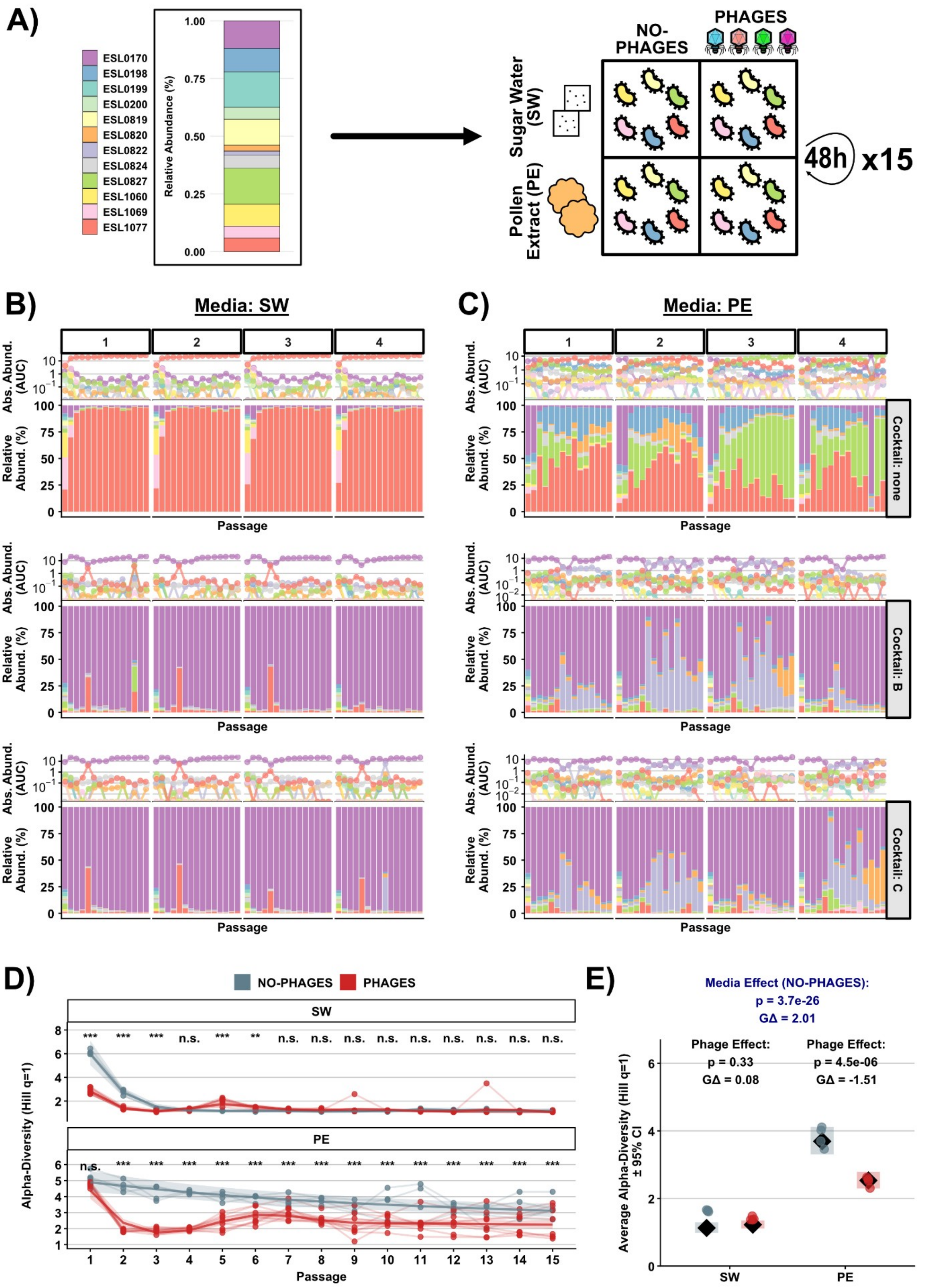


**Fig. S6: Phage effect in the 12-strain pilot community is dampened by a phage-resistant strain.**

(**A**) Experimental design. All 12 *Bifidobacterium* strains were inoculated at equal proportions (left) and grown in a full-factorial longitudinal experiment across two resource conditions (sucrose water, SW; pollen extract, PE) and three phage treatments (NO-PHAGES, Cocktail B, Cocktail C), with four technical replicates per condition, serially transferred every 48h for 15 passages. (**B-C**) Absolute abundances (top, AUC) and relative abundances (bottom, stacked bar plot) of all 12 strains across 15 passages in SW (B) and PE (C), for NO-PHAGES control (top rows), Cocktail B (middle rows), and Cocktail C (bottom rows). Each column of panels corresponds to one replicate. Strains are color-coded as in (A). (**D**) Alpha diversity (Hill q=1) over time in SW (top) and PE (bottom), in the absence or presence of phages. Connected points show individual replicate trajectories. Lines show GAM-predicted marginal means ± 95% CI. Asterisks denote significant pairwise differences between phage treatments at each passage (n.s., not significant; ** $p < 0.01$; *** $p < 0.001$). (**E**) Average alpha diversity across passages summarized by condition. Diamonds show estimated marginal means from GAM. Crossbars show 95% CI. Points show replicate averages. Statistics indicate the effect of media (NO-PHAGES only, blue) and the effect of phages within each media condition (GΔ, Glass's Δ). All p-values are corrected for multiple comparisons using the FDR method.

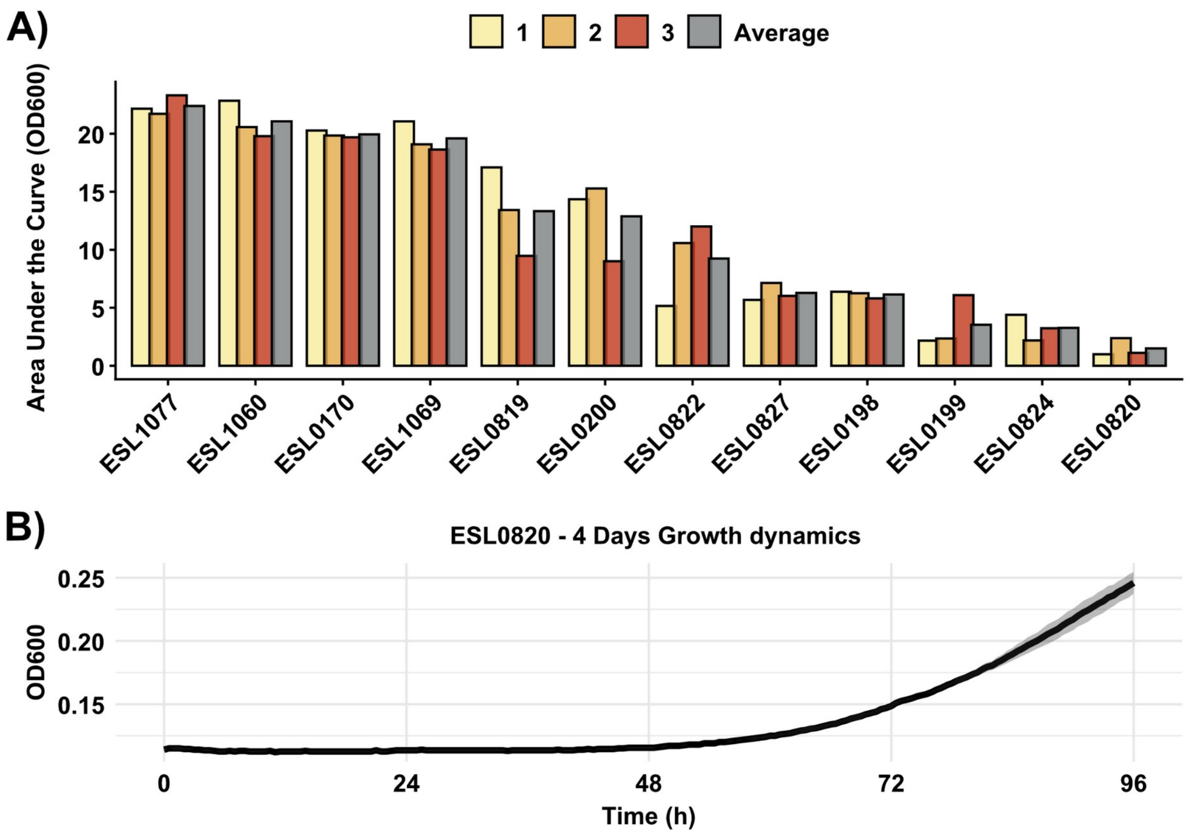


**Fig. S7: Monoculture growth of *Bifidobacterium* strains in sucrose water.**
(**A**) Monoculture area under the growth curve (AUC, OD600) in sucrose water of the 12 *Bifidobacterium* strains used in this study across three passages (fig. S2) and their average (grey). Strains are ordered by decreasing average AUC. (**B**) Growth curve of ESL0820 measured over 96h in monoculture.

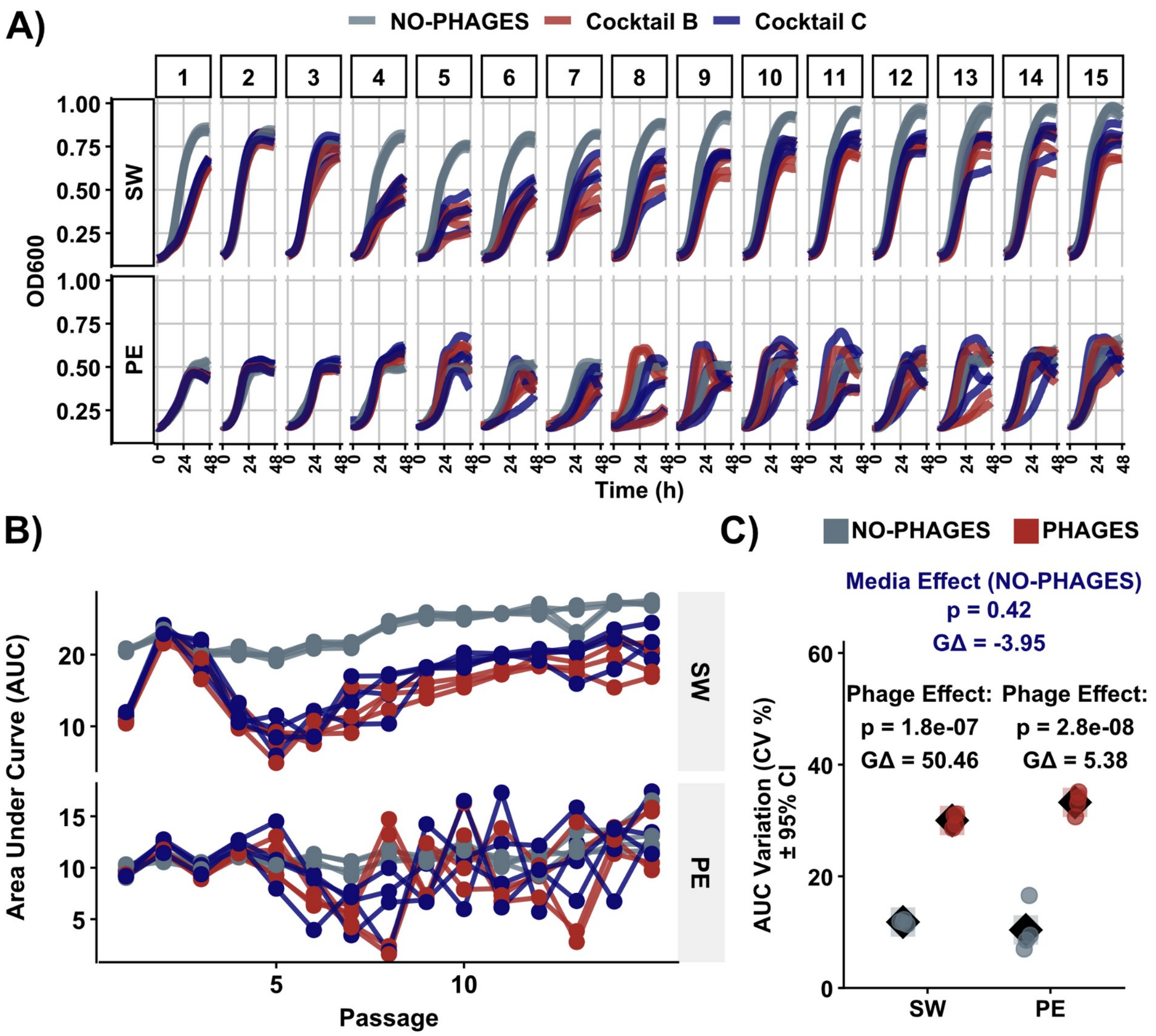


**Fig S8: Phage effect on community growth in pilot experiment with a 12-strains community.**

(**A**) Growth curves (OD600 over 48h) of a pilot 12-strain community across 15 successive passages in SW (top) and PE (bottom), in the absence of phages (grey) or in the presence of enriched phage cocktail B or C (red). Each line represents the growth dynamics of one technical replicate. (**B**) Community productivity (area under the growth curve, AUC) over passages for each condition and media. Connected points show individual replicate trajectories. (**C**) Temporal variability in community AUC, measured as the coefficient of variation (CV%) across passages. Diamonds show estimated marginal means. Crossbars show 95% CI. Points show replicate values. Statistics indicate the effect of media (NO-PHAGES only) and the effect of phages within each media condition (GΔ, Glass's Δ). All p-values are adjusted for multiple comparisons using the FDR method.

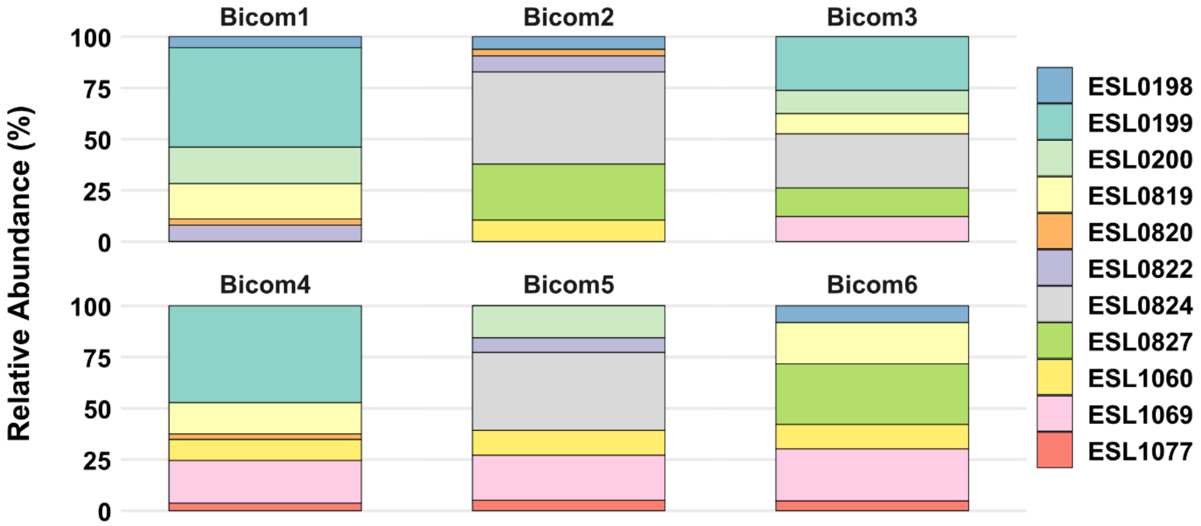


**Fig. S9: Initial composition of the six synthetic *Bifidobacterium* communities.** Relative abundance of each *Bifidobacterium* strain at inoculation across the six communities (Bicom1-6). Each community contains six strains.

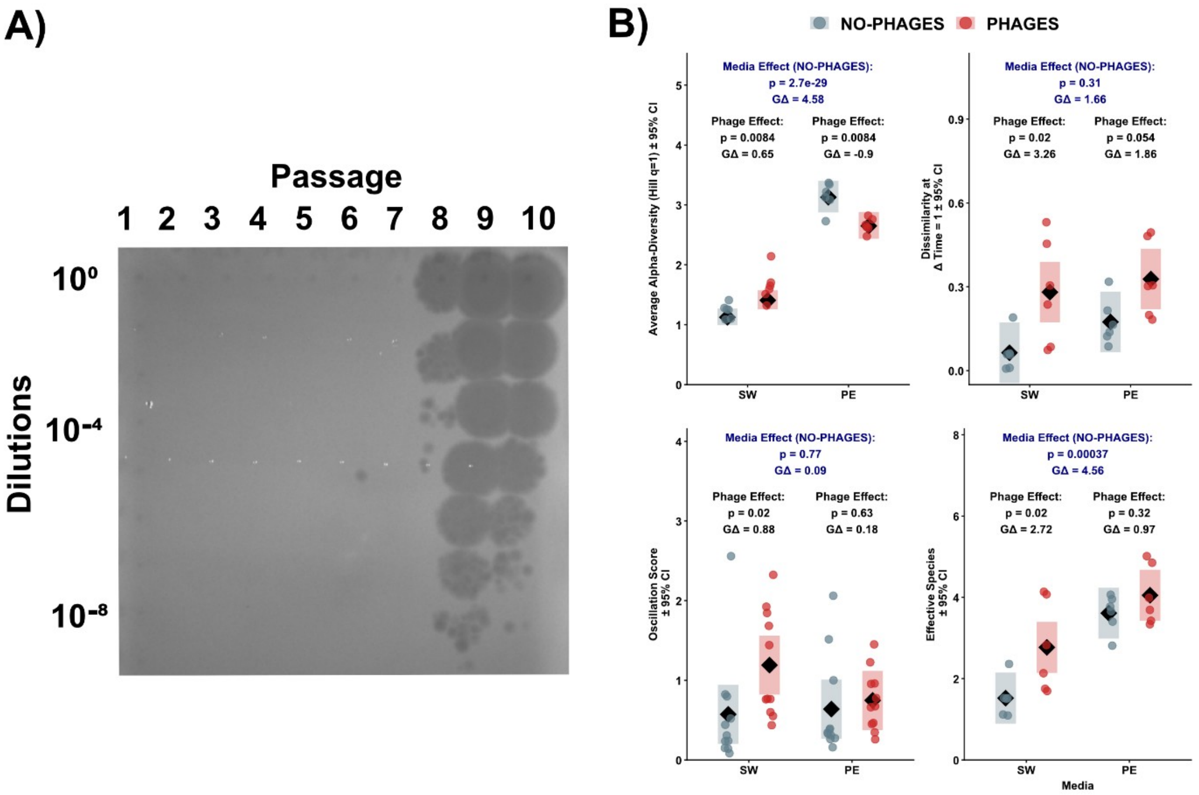


**Fig. S10: Phage contamination in the no-phage sample of Bicom2 on PE and analysis of its effect on the overall results.**

(**A**) Photo of plaques obtained from the culture of Bicom2 on PE without phages across passages 1-10 at serial dilutions ($10^0$ to $10^{-8}$). Plaques are visible from passage 8 onward. (**B**) Sensitivity analysis excluding data from passages 8-10 of the Bicom2 culture on PE without phages. All four diversity metrics are shown: average alpha diversity (Hill q=1, top left), short-term community turnover (Bray-Curtis dissimilarity at Δtime = 1, top right), oscillation score (bottom left), and temporal gamma diversity (effective species, bottom right). Diamonds show estimated marginal means. Crossbars show 95% CI. Points show community-level values. Statistics indicate the effect of media (NO-PHAGES only) and the effect of phages within each media condition (GΔ, Glass's Δ). All results are qualitatively unchanged relative to the full dataset, confirming that the contamination did not affect our conclusions.

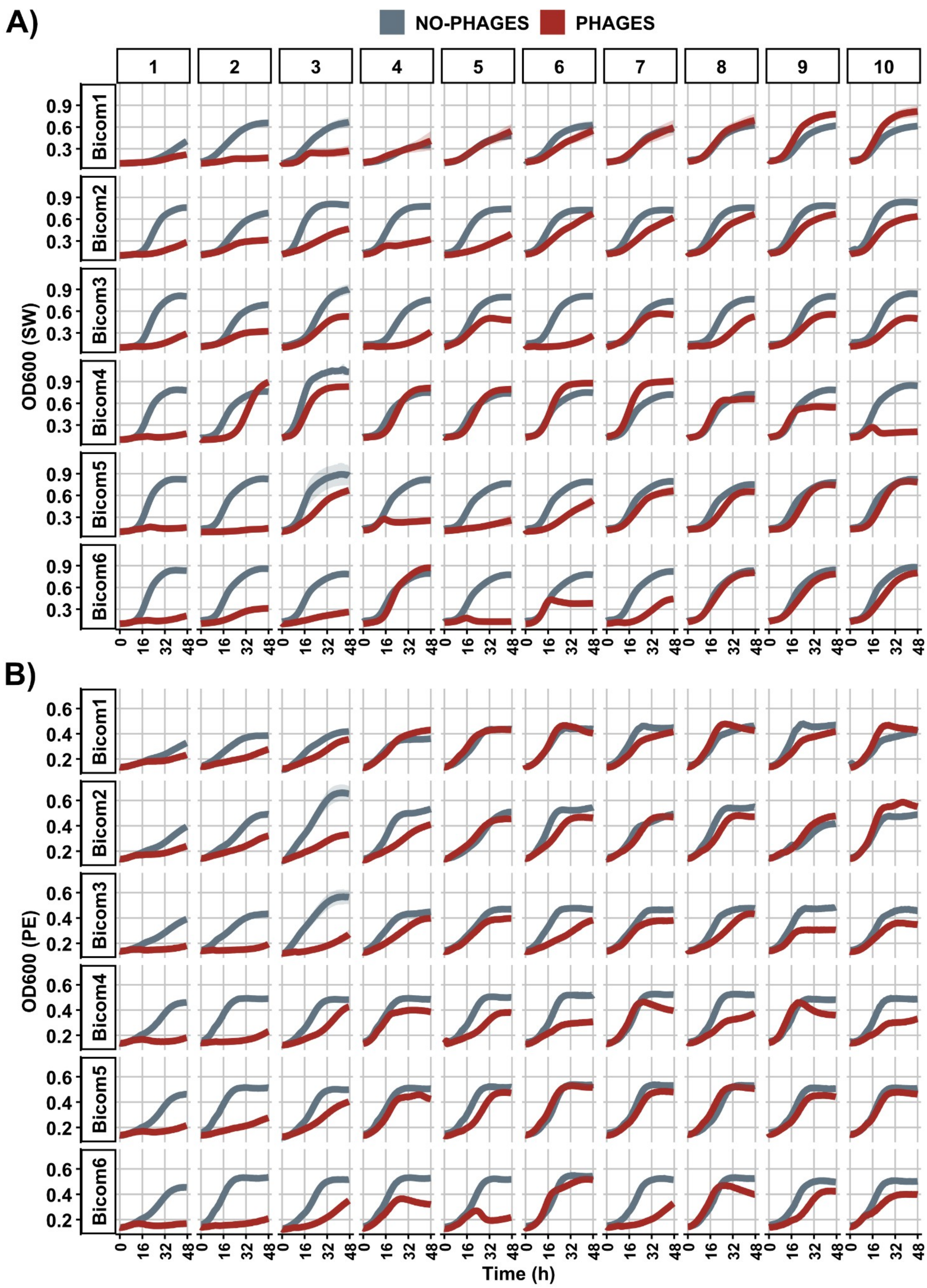


**Fig. S11: Community-level growth curves across passages in the main experiment.** OD600 measured over 48h for each of the six synthetic communities (Bicom1-6, rows) across 10 passages (columns), in the absence or presence of phages, in (**A**) sucrose water (SW) and (**B**) pollen extract (PE). Lines show the mean across replicates. Shaded ribbons show the standard deviation over three technical replicates.

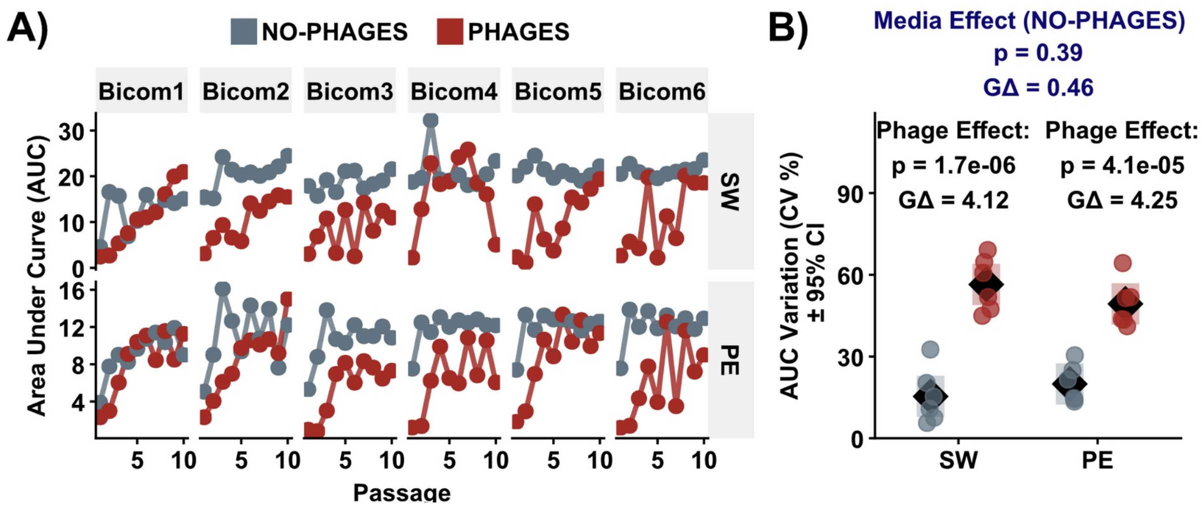


**Fig. S12: Effect of phage predation on community growth.**

(**A**) Community-level area under the growth curve (AUC) across 10 passages for each of the six communities (Bicom1-6) in sucrose water (SW, top) and pollen extract (PE, bottom), in the absence or presence of phages. Connected points show individual replicate trajectories. (**B**) Temporal variability in community AUC, measured as the coefficient of variation (CV%) of AUC across all passages. Diamonds show estimated marginal means. Crossbars show 95% CI. Points show community-level average values. Statistics indicate the effect of media (NO-PHAGES only) and the effect of phages within each media condition (GΔ, Glass's Δ).

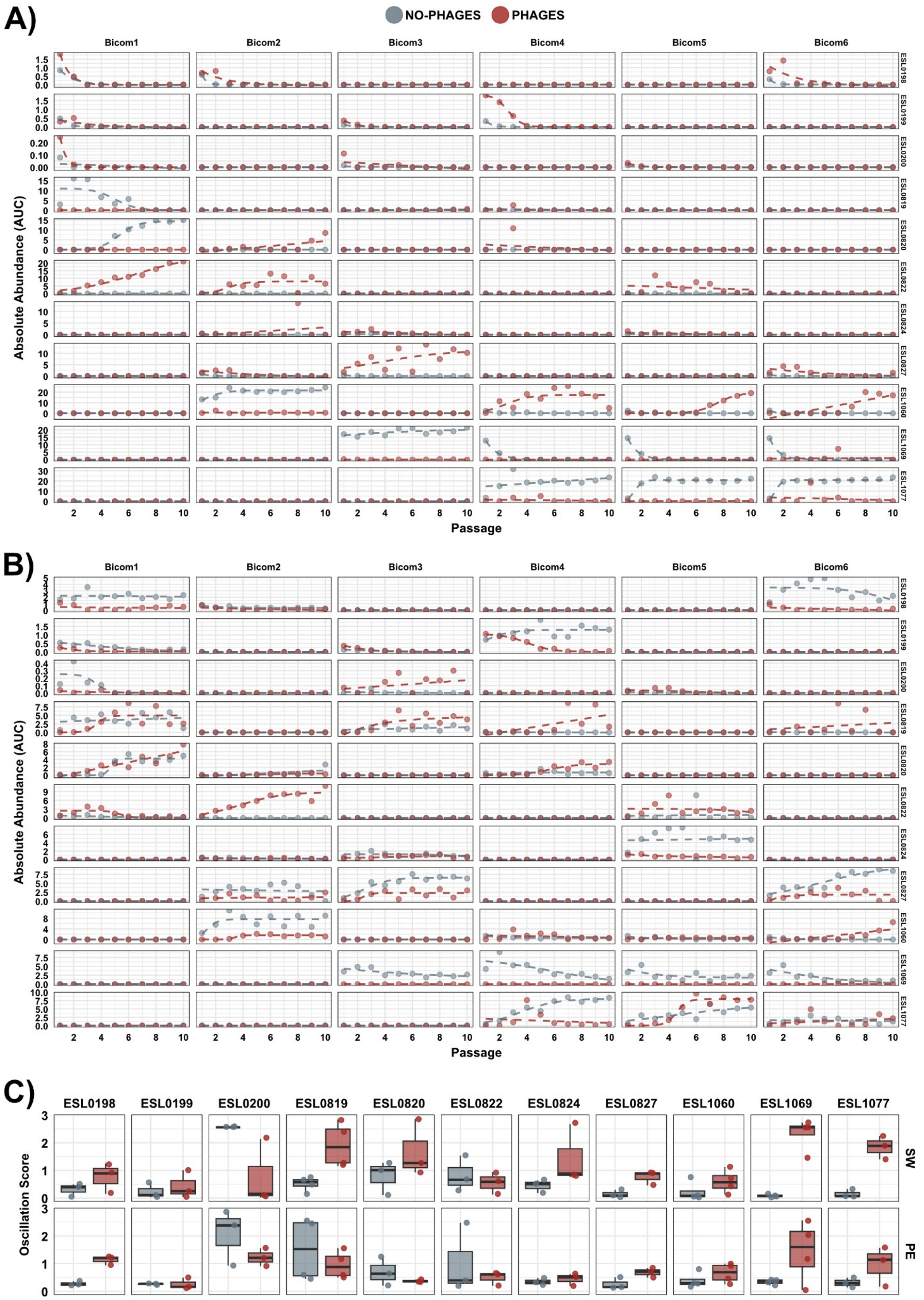


**Fig. S13: Strain-level absolute abundance trajectories and oscillation scores across phage and resource conditions.**

(**A-B**) Absolute abundance (AUC) of each strain (rows) across 10 passages in each of the six communities (columns), in (A) sucrose water (SW) and (B) pollen extract (PE), in the absence (grey) or presence (red) of phages. Points show measurements. Dashed lines show non-oscillating growth model fit. (C) Boxplots of oscillation scores (CV_RMSE) for each strain in SW (top) and PE (bottom), in the absence or presence of phages. Each panel shows the distribution across communities in which the strain is present. Boxes show the interquartile range; horizontal lines show the median; points show individual community values.

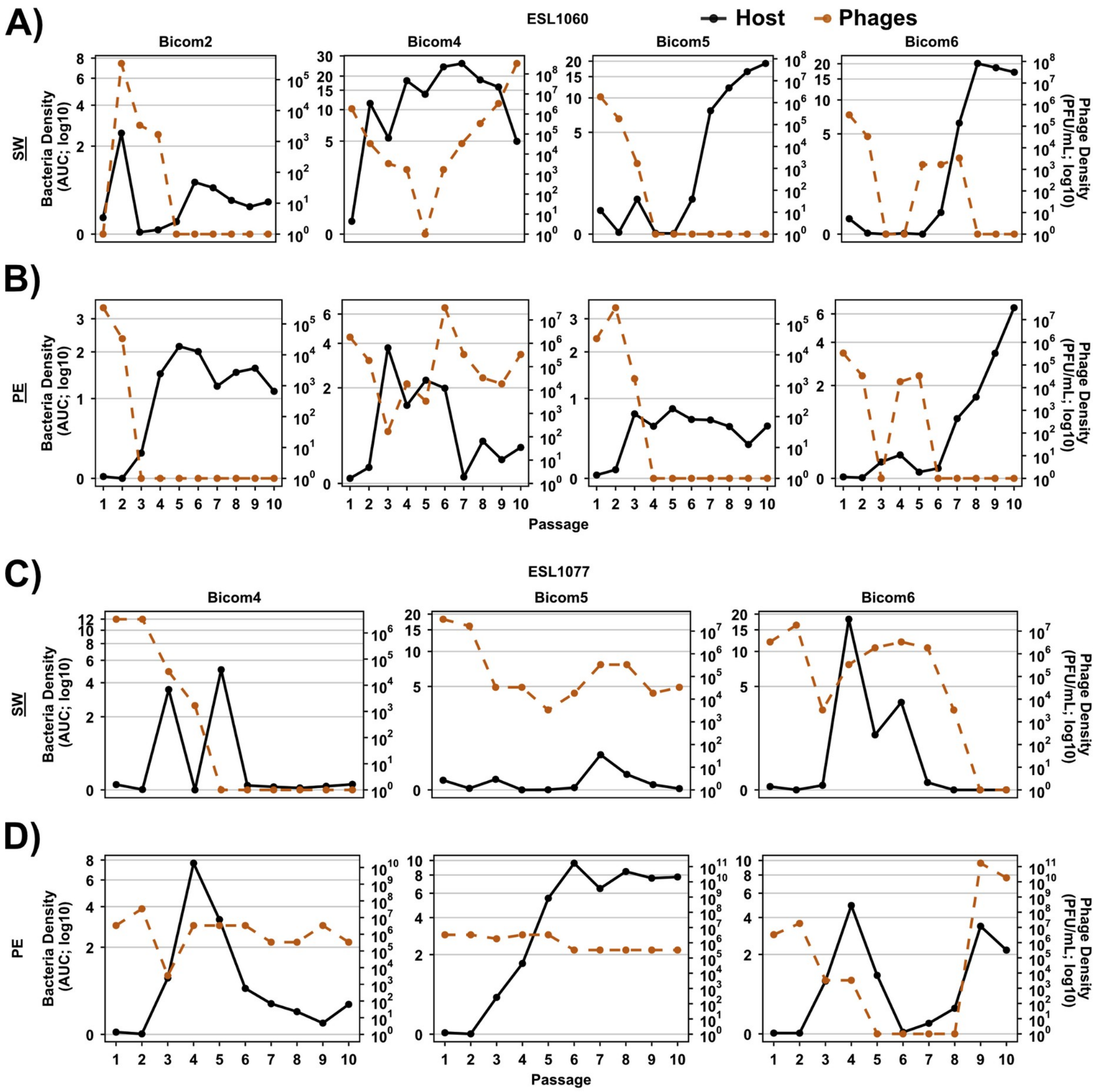


**Fig. S14: Longitudinal tracking of phage and host densities for ESL1060 and ESL1077.** Absolute bacterial abundance (AUC, log10; black solid line, left y-axis) and phage density (PFU/mL, log10; orange dashed line, right y-axis) tracked over 10 passages for strain (**A-B**) ESL1060 and (**C-D**) ESL1077, in communities where each strain is present, across (**A, C**) SW and (**B, D**) PE conditions. Each panel corresponds to one community (Bicom identity indicated at top).

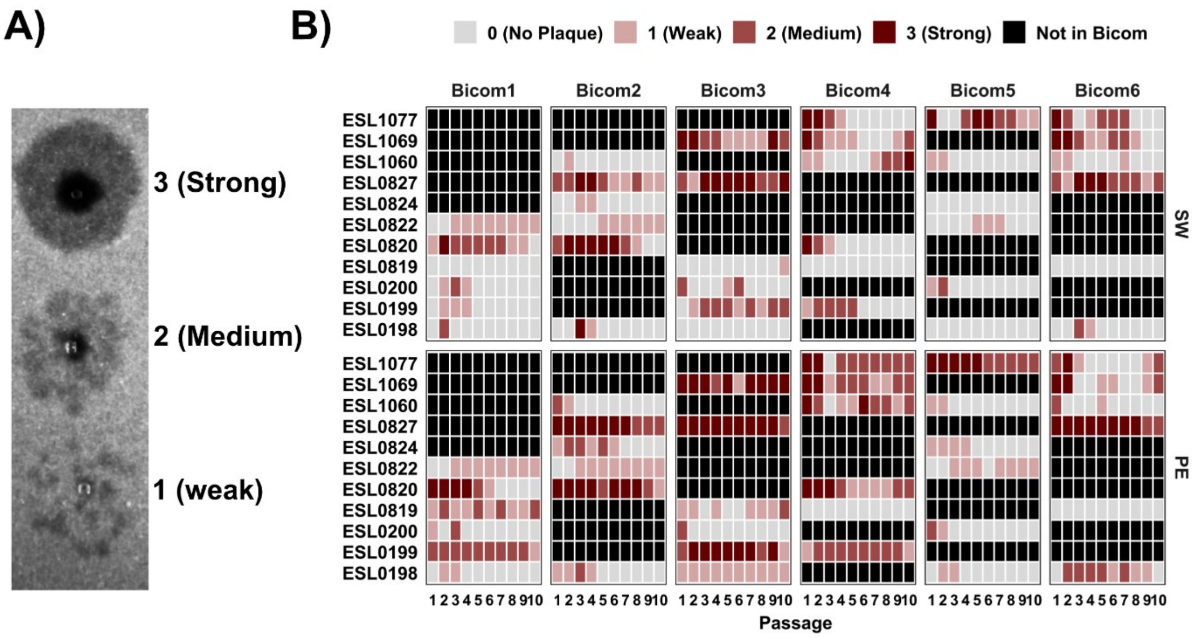


**Fig. S15: Longitudinal infectivity profiles of phage communities across passages.**
**(A)** Plaque formation was scored on a qualitative scale from 0 (no plaque) to 3 (strong plaque). **(B)** Infectivity profiles were assessed by spot assay for each bacterial strain (rows) across 10 passages (columns), and for each of the six communities (Bicom1-6) in SW (top) and PE (bottom). Black cells indicate strains absent from a given community. Each cell represents the score from a spot assay of the phage community collected at that passage against the indicated bacterial strain. The full matrix of scores across communities and passages constitutes the infectivity profile used to quantify plaque richness and temporal turnover in Fig. 4.

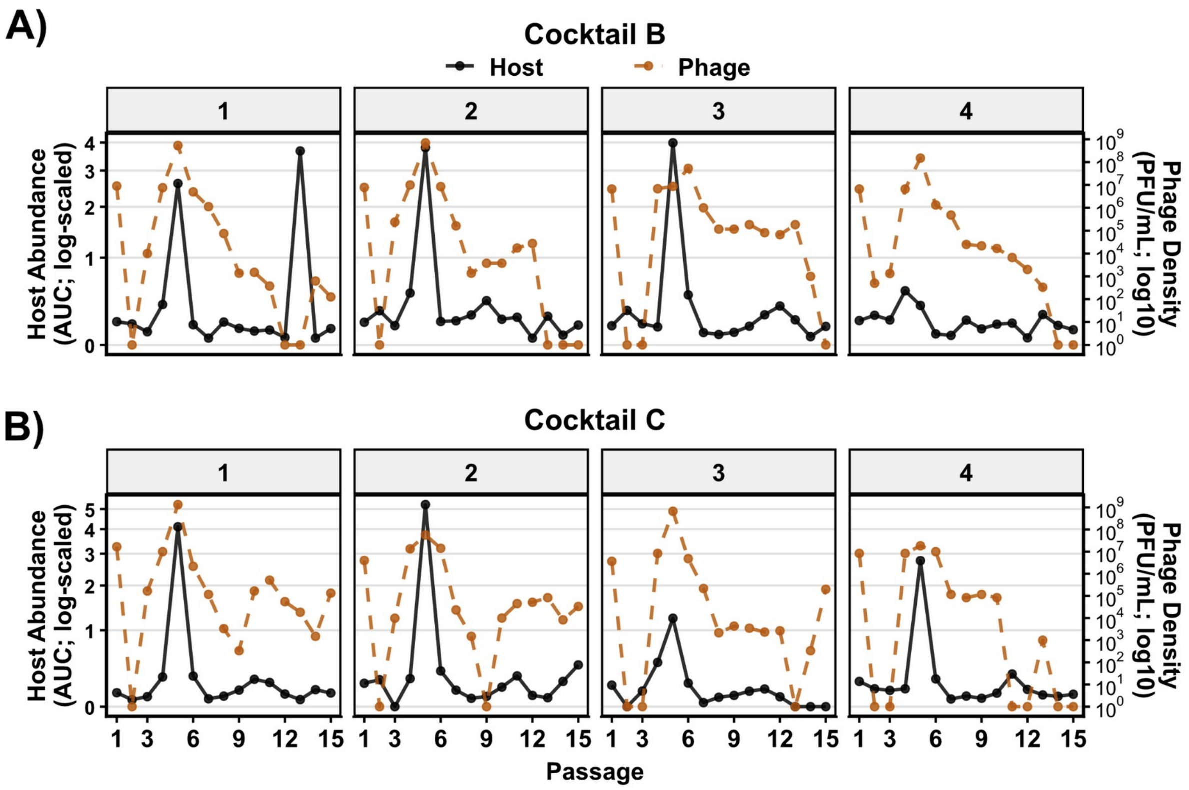


**Fig. S16: Predator-prey dynamics in the pilot 12-strain community.**

Absolute host abundance (AUC, log-scaled; black solid line, left y-axis) and phage density (PFU/mL, log10; orange dashed line, right y-axis) tracked across 15 passages for ESL1077 in the pilot experiment, with (**A**) cocktail B and (**B**) cocktail C. Each panel corresponds to one of four technical replicates. In both cocktail conditions, bacterial abundance peaks are closely followed by increases in phage density, consistent with predator-prey dynamics. This pattern is considerably clearer than in the 6-strain communities (Fig. S14), likely reflecting the near-two-strain nature of this system, where community dynamics are effectively governed by the dominant susceptible strain (ESL1077) and the phage-resistant strain (ESL0170).

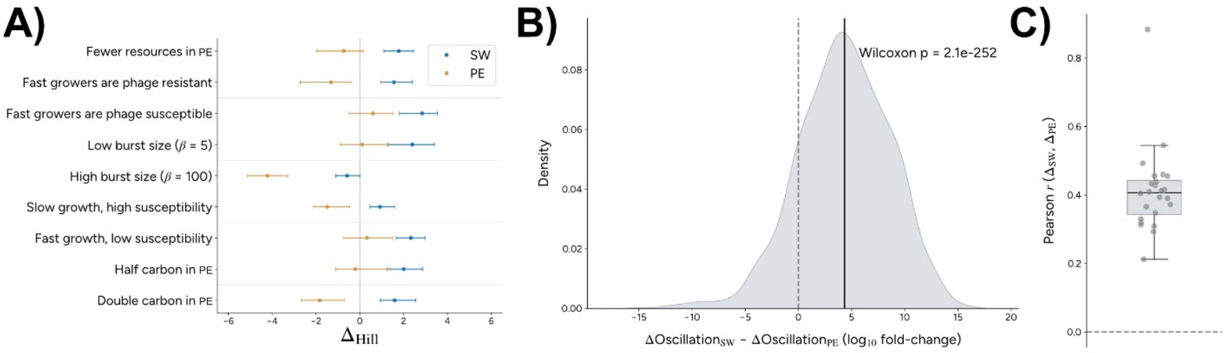


**Fig. S17: Robustness of mathematical model results across parameter variations.**
(**A**) Median change in alpha -diversity (Hill q=1) due to phages (ΔHill = difference in alpha-diversity between PHAGES and NO-PHAGES conditions) in sucrose water (SW, blue) and pollen extract (PE, orange) across different conditions of resource number, growth-defense tradeoff, burst size, trait scales, and total carbon supply. Error bars show the interquartile range across 100 simulated communities. In all configurations, ΔHill is larger in SW than in PE, confirming that niche overlap, not any specific parameter choice, drives the media-dependent phage effect. (**B**) Distribution of comparison of phage-induced oscillations (ΔOscillation_SW - ΔOscillation_PE), pooled across all parameter configurations. Phages induce stronger oscillations in SW than in PE (Wilcoxon signed-rank test, $p < 0.001$). Solid black line represents the median difference in oscillation score, and gray dashed line marks zero. (**C**) Pearson correlation between ΔHill in sucrose water and pollen extract, computed across 24 parameter configurations spanning the 9 scenarios shown in panel A. All conditions were significant ($p<0.05$, N=100 communities tested per configuration).

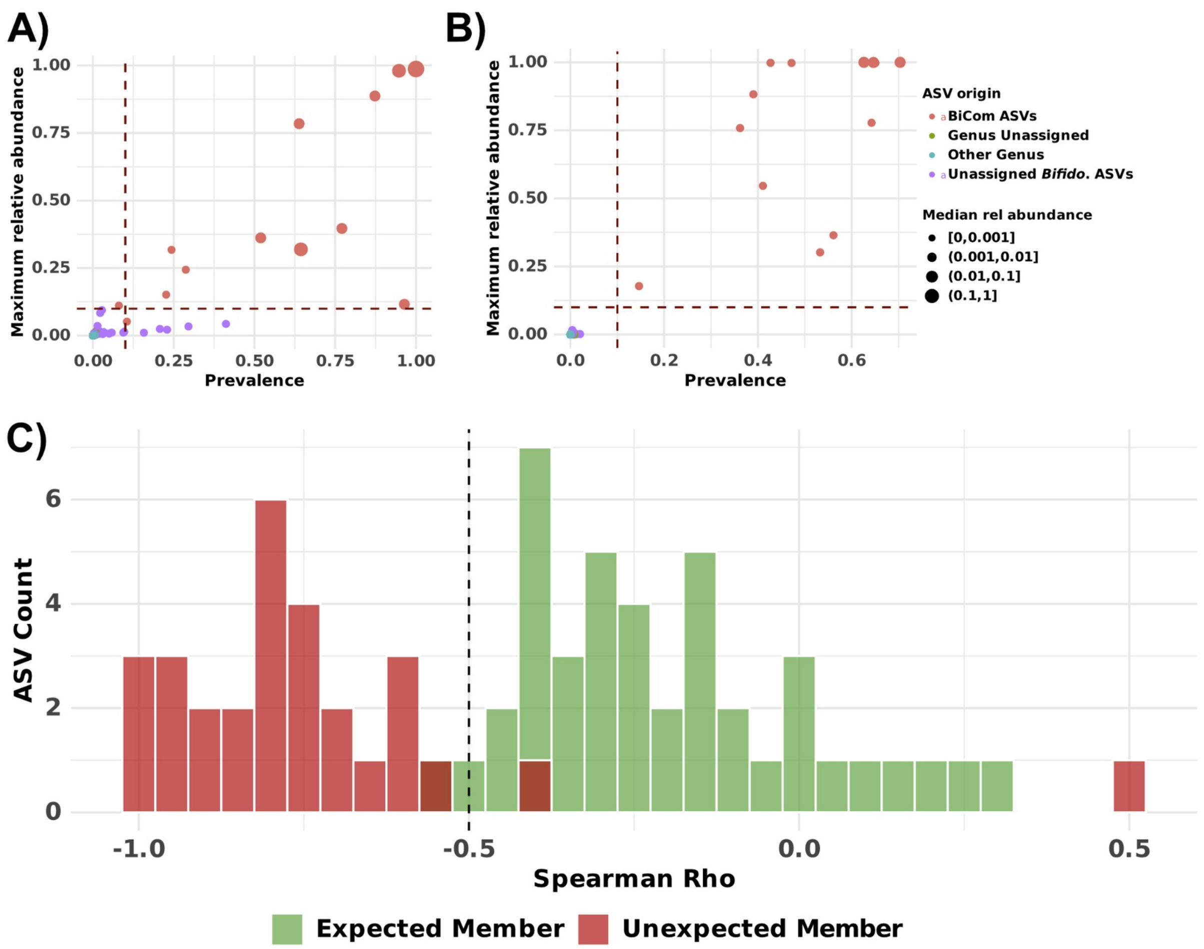


**Fig. S18: Quality filtering of ASVs in the passaging experiments.**

(**A**) Prevalence versus maximum relative abundance of all ASVs detected in the 12-strain passaging experiment, colored by ASV origin. BiCom ASVs (pink) correspond to strains included in the synthetic community; Unassigned Bifidobacterium ASVs (purple) correspond to reads assigned to the genus but not to any included strain; Other Genus (teal) and Genus Unassigned (yellow) correspond to non-Bifidobacterium ASVs. Point size reflects median relative abundance. Dashed lines indicate the prevalence (10%) and maximum relative abundance (10%) thresholds used for filtering. ASVs falling below both thresholds were removed. (**B**) Same plot for the 6-strain passaging experiment. (**C**) Distribution of Spearman correlation coefficients between the relative abundance of unexpected ASVs and library size in the 6-strain experiment, colored by whether the ASV corresponds to an expected (green) or unexpected (red) community member. Unexpected ASVs showed strongly negative correlations with library size ($\rho < -0.5$, dashed line), consistent with cross-well contamination during DNA extraction or library preparation rather than genuine biological presence and were removed from downstream analyses.
